# A transcriptional mechanism involving R-loop, m6A modification and RNA abasic sites regulates an enhancer RNA of *APOE*

**DOI:** 10.1101/2022.05.01.489793

**Authors:** Jason A. Watts, Christopher Grunseich, Yesenia Rodriguez, Yaojuan Liu, Dongjun Li, Joshua T Burdick, Alan Bruzel, Robert J. Crouch, Robert W. Mahley, Samuel H. Wilson, Vivian G. Cheung

## Abstract

The DNA genetic code and the RNA regulatory code determine phenotypes from gene expression to disease susceptibility. DNA sequence variants lead to phenotypic differences among individuals, while within an individual, RNA dynamically confers cell identity and responds to cellular and environmental signals. To provide regulation for different cell types and conditions, the nucleotides of RNA are modified by hundreds of chemical reactions, and RNA folds into innumerable shapes. To decipher the RNA regulatory code is to understand how RNA sequence and structure respond to cellular needs. Here, we deciphered one part of the RNA code where RNA abasic sites in R-loops regulate transcription by pausing RNA polymerase II. We uncovered an enhancer RNA, AANCR, that regulates the transcription and expression of *APOE*. When AANCR is folded into an R-loop, which is then modified by N6-adenine methylation and N-glycosidic cleavage, it is a partially transcribed nonfunctional enhancer and *APOE* is not expressed. In contrast, in some cell types and under stress, AANCR does not form a stable R-loop as its sequence is not modified, so it is transcribed into a full-length enhancer that promotes *APOE* expression. By genetic analysis, we confirmed that AANCR regulates *APOE* expression. DNA sequence variants in AANCR are associated with *APOE* expression and also with Alzheimer’s disease. Our data show that DNA and RNA sequence and structure jointly regulate gene expression that influence disease risk.

**Highlights:**

- m6A, RNA abasic sites and R-loops jointly regulate transcription by pausing RNA Polymerase II.
- An enhancer RNA regulates *APOE* expression.
- Enhancer RNA of *APOE* modifies susceptibility to Alzheimer’s disease.

## Introduction

DNA and RNA constitute the genetic and regulatory code of all organisms. DNA sequence differences account for individual variation from gene expression (Cheung et al., 2005; Morley et al., 2004) to disease susceptibility. While technological advances have made it easier to identify DNA sequence mutations and variants that account for the genetic bases of diseases, molecular understanding remains a challenge. To gain deeper insight, it is necessary to delve into both the genetic and the regulatory codes. RNA sequence and structure comprise the regulatory code. RNA is composed of four canonical bases (A, C, G, and U) and over 150 modified bases (Boccaletto et al., 2018) that form a myriad of structures. In an organism, every cell has largely the same DNA, but different RNA species. RNA is what confers cell identity and function. It is templated from DNA, but during and after synthesis, it is highly processed by hundreds of chemical steps that modify the bases and sugar of RNA. Chemical modifications of RNA include methylation that generates N6-methyladenosine to multiple-step reactions that form wybutosine.

RNA sequence and structure are closely related, they regulate each other and co-regulate gene expression and function. RNA sequence affects its structure and conversely, its structure can further alter the sequence, as in RNA editing in which (adenosine-uridine) AU-rich sequences form stem-loop structures that are bound by ADAR proteins to deaminate adenosine to inosine (Mannion et al., 2015; Stefl et al., 2010). RNA sequence and structure also mediate the interaction between RNA and regulatory nucleic acids and proteins; for example, the repression of mRNA by microRNA and Argonaut-2 is dependent on both sequence and structure (Schirle et al., 2014).

Our study of amyotrophic lateral sclerosis type 4, a juvenile-onset ALS due to heterozygous senataxin mutation showed that patients’ cells have significantly fewer R-loops (Grunseich et al., 2018, 2020), three-stranded nucleic acid structures, each with an RNA/DNA hybrid and a displaced singlestranded DNA (Aguilera and Garcia-Muse, 2012; Castillo-Guzman and Chédin, 2021; Niehrs and Luke, 2020; Thomas et al., 1976). To understand how the deficiency of R-loops affects cell function, we first looked for proteins that bind to these nucleic acid structures. We and other groups have identified hundreds of R-loop binding proteins (Cristini et al., 2018; Wang et al., 2018; Yan et al., 2022). These studies use different methods, yet the results consistently show the same set of several hundred proteins. These include a significant number of enzymes that modify nucleic acids such as: METTL3 and METTL14 that methylate adenosine in RNA to form N6-methyladenosine, as well as methylpurine glycosylase (MPG) and apurinic/apyrimidinic endonuclease 1 (APE1) that were known to process DNA (Lindahl, 1974). These results suggest that R-loops serve as platforms for processing nucleic acid including RNA modification. Yet to confirm their regulatory roles, the mechanism must be determined for how each protein processes nucleic acid individually and jointly.

We first focused on two proteins, MPG and APE1, and found that MPG not only cleaves the N-glycosidic bond on DNA but also on RNA, leading to RNA abasic sites (Liu et al., 2020). APE1 then processes the RNA by cleaving the sugar-phosphate backbone at the abasic sites. The activity of MPG and APE1 on RNA occurs only when the RNA is hybridized to a DNA strand, as in an R-loop, thus further illustrating the co-dependence of sequence and structure.

In the current study, we set out to examine RNA abasic sites in R-loops and uncovered a gene regulatory mechanism by which an RNA abasic site stabilizes an R-loop to pause RNA Polymerase II transcription. We found an enhancer RNA of *APOE* whose expression and function are regulated dynamically by pausing of RNA Polymerase II. When the noncoding RNA is full-length, it activates *APOE* expression. When the enhancer RNA is only partially transcribed, it is nonfunctional and *APOE* is not expressed, because RNA Polymerase II transcription is paused by R-loops that are stabilized by RNA abasic sites that form when MPG cleaves N6-methyladenines. In response to stress, the R-loops that pause transcription are resolved, the noncoding RNA is transcribed into a full-length enhancer that activates *APOE* expression. By genetic analysis, we showed that sequence variants in the enhancer region affect *APOE* expression, including in the hippocampus, and are associated with Alzheimer’s Disease. Thus, we have revealed a gene regulatory mechanism in which the sequence and structure of a noncoding RNA regulate its transcription, function and in turn disease susceptibility.

## Results

### Methylpurine glycosylase knockdown induces *APOE* expression

In skin fibroblasts, we knocked down MPG by RNA interference and carried out RNA sequencing which showed that the most highly induced gene is *APOE*. *APOE* is not expressed or at a very low level in skin fibroblasts, but when MPG is knocked down, *APOE* gene expression is highly induced (Figure 1A, >50-fold). The immunoblot shows that APOE protein expression is also upregulated significantly (Figure 1B, P<<0.001, >20-fold). The resultant APOE is then secreted, as APOE protein level is significantly higher (Figure 1C, P<<0.001, >50-fold) in the cell culture media for the fibroblasts whose MPG is knocked down. We also knocked down MPG in lung epithelial cells and found that APOE protein expression is also induced in these cells (Supplemental Figure 1). We then carried out PRO-seq and found that upon MPG knockdown, there is an increase in the abundance of RNA Polymerase II (RNA Pol II) in the *APOE* promoter and active transcription of *APOE* (Figure 1D). Thus, MPG knockdown induces transcription and expression of *APOE*.

**Figure 1.**
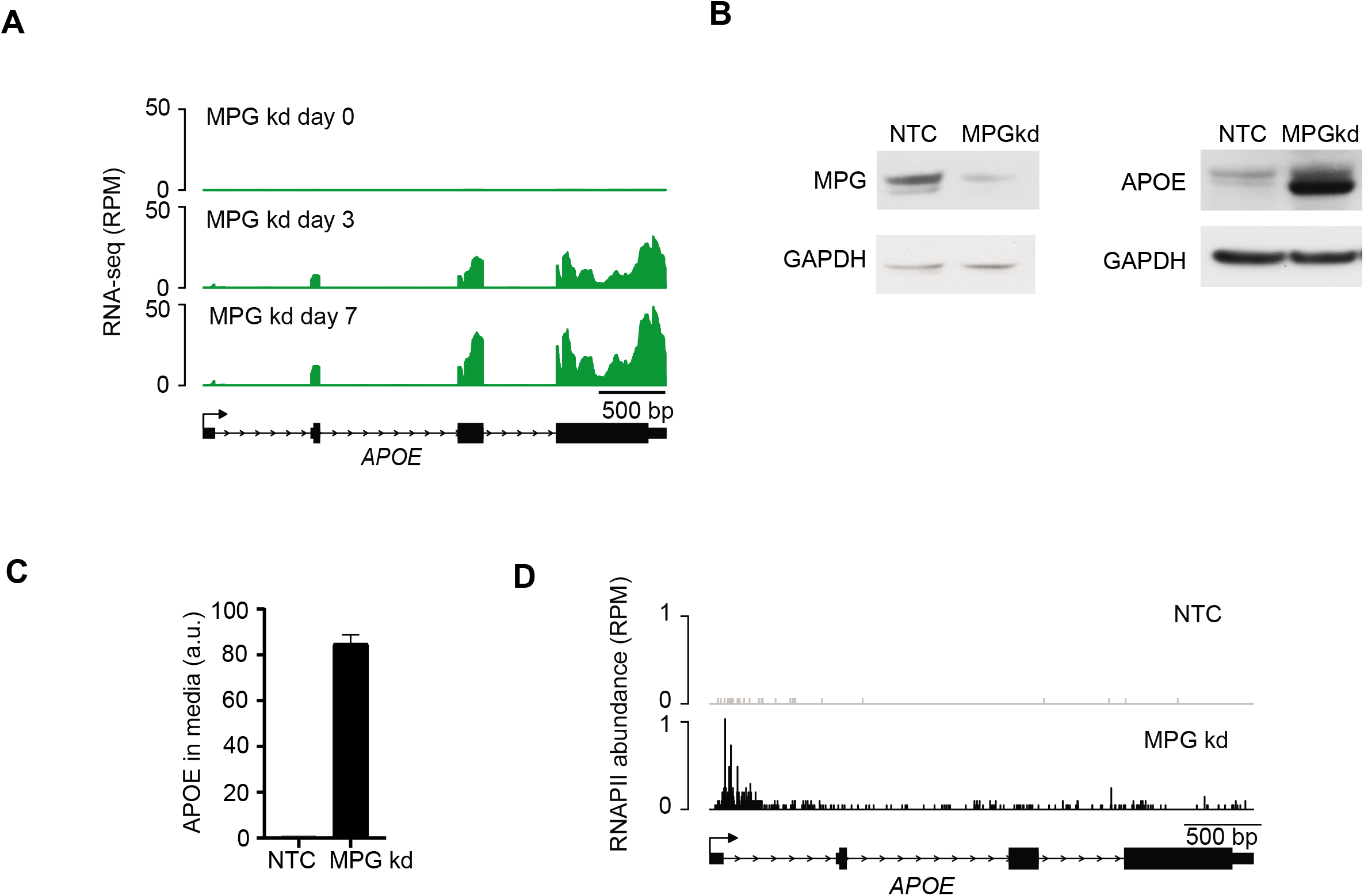
APOE is regulated by MPG. (A) Representative RNA-seq showing *APOE* expression in fibroblasts before and after MPG knockdown (days 3 and 7), Y-axis is RPM. (N=2), (B) Representative immunoblot of MPG and APOE expression before and after MPG knockdown (day 7). GAPDH is a loading control. (N>5), (C) Densitometry quantification of immunoblots of secreted APOE protein before and after MPG knockdown (day 7) (N=3, P<<0.001 t-test, error bars=S.E.M.). NTC is the non-target control. (D) Representative PRO-seq results for *APOE* before and after MPG knockdown (day 7) are plotted. The bar at each nucleotide location represents the number of RNA Pol II. *APOE* transcription is induced after MPG knockdown. Y-axis is RPM (N=2).

### R-loops form upstream of *APOE*

Our previous study showed that MPG forms abasic sites in the RNA of RNA-DNA hybrids (Liu et al., 2020). Here, we asked whether MPG affects *APOE* expression through binding to R-loops. Using the S9.6 antibody that specifically recognizes R-loops (Hu et al., 2006; Kosar et al., 2021), we carried out DNA-RNA immunoprecipitation, DRIP, followed by sequencing. Since *APOE* is not transcribed or at a very low level in fibroblasts, there was no RNA to form R-loops in *APOE*, rather we detected an R-loop upstream of *APOE* (Figure 2A, top panel), in an intergenic region. This upstream region is transcribed by RNA Pol II until it pauses just 5’ to the R-loops (Figure 2A, bottom panel). To assess if the R-loops or NELF protein complex pauses the RNA Pol II transcription (Yamaguchi et al., 1999), we carried out immunoprecipitation against one of the NELF proteins, NELFA. We did not detect NELFA binding to the region upstream of *APOE*, whereas NELFA binding was readily detected at the *HSP70* promoter (Figure 2B). This suggests that in the region upstream of *APOE*, RNA Pol II transcription is paused by R-loops.

**Figure 2.**
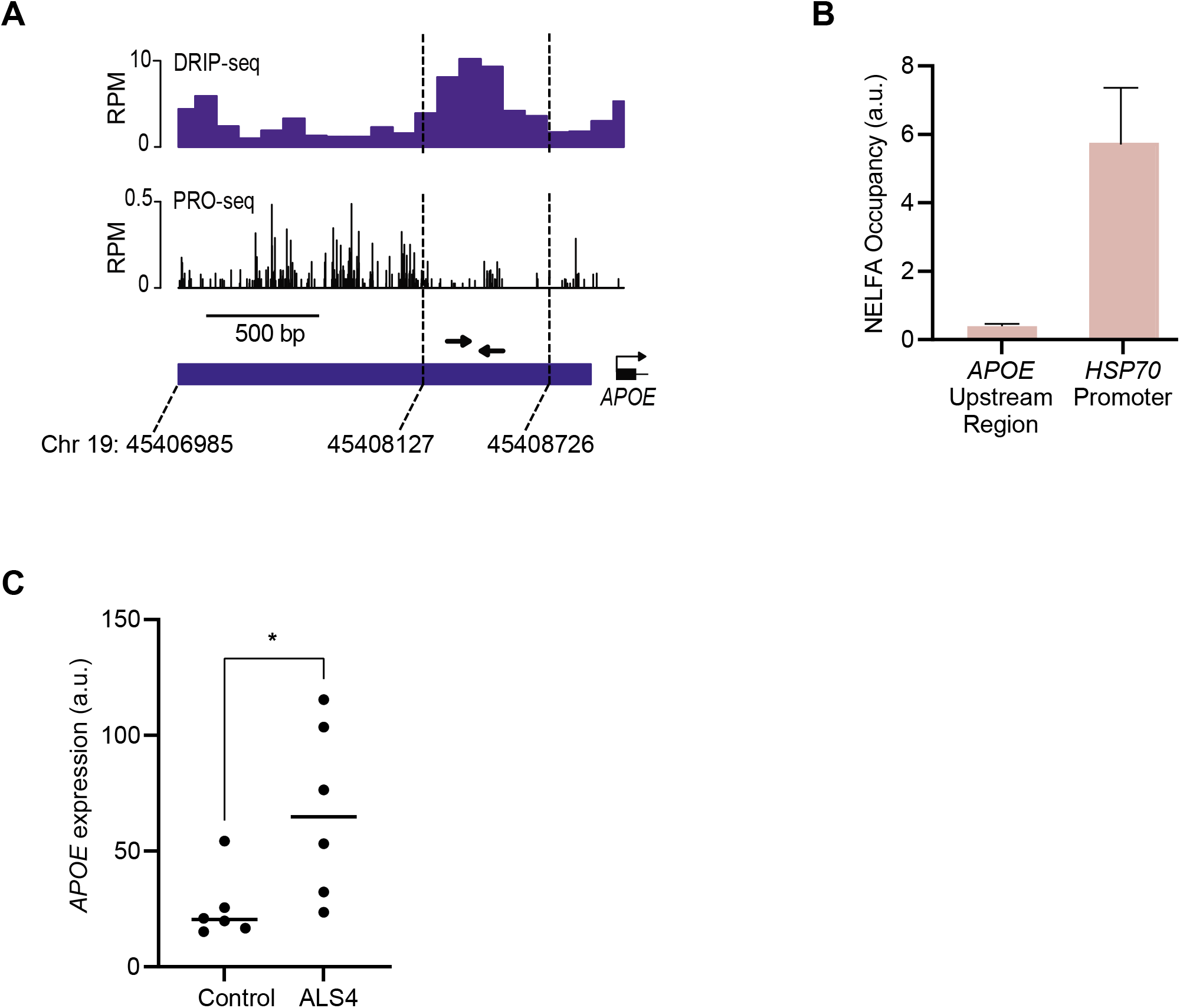
Upstream R-loops regulate APOE expression. (A) Average S9.6 DRIP-seq data in fibroblasts (N=5) are plotted (top panel) and show R-loops upstream (5’) of *APOE*. Average PRO-seq results in fibroblasts (N=5) are plotted (bottom panel), showing active transcription in the intergenic region upstream of *APOE*, and RNA Pol II pausing upstream of the R-loops. Y-axis is RPM. Arrows indicate qPCR primers used in subsequent figures. Genomic coordinates (hg19) are indicated. (B) Data from NELFA chromatin immunoprecipitation followed by qPCR with primers indicated by arrows in Figure 2A are shown as fold-enrichment over IgG in arbitrary units (a.u.). (N=2, error bars are S.E.M). (C) *APOE* expression levels in fibroblasts from family-control and ALS4 patients are plotted, each dot represents *APOE* expression of one person, (*P<0.05, t-test).

We wondered if, in this region, the R-loops affect *APOE* expression. We examined *APOE* expression in ALS4 patient cells. ALS4 encodes a hyperactive senataxin (RNA/DNA helicase) that reduces R-loops without affecting transcription initiation (Chen et al., 2004; Grunseich et al., 2018). We compared *APOE* expression in fibroblasts from ALS4 and controls. We found that on average, ALS4 patients with fewer R-loops (Grunseich et al., 2018) have higher *APOE* expression (Figure 2C). While *APOE* is not expressed in most of the fibroblasts from controls, it is expressed in many of the fibroblasts from ALS4 patients. Thus, cells with fewer stable R-loops have higher *APOE* expression. Both MPG expression and the R-loop abundance are negatively correlated with *APOE* expression.

### MPG generates RNA abasic sites in the R-loops upstream of *APOE*

Next, we asked if the MPG binds to the R-loops near *APOE* and if so, are RNA abasic sites generated. We carried out MPG RNA-IP and ARP RNA-pulldown. Aldehyde-reactive probe (ARP) is a biotinylated amine that binds to the aldehyde on the exposed sugar of an abasic site (Kubo et al., 1992; Tanaka et al., 2011), thus ARP RNA-pulldown identifies abasic site-containing RNA. We confirmed by S9.6 DRIP-PCR (Figure 3A, left panel), the R-loops upstream of *APOE* as shown in Figure 2A. MPG RNA-IP then shows that MPG binds to the RNA in those R-loops upstream of *APOE* (Figure 3A, middle panel), and ARP RNA-pulldown (Figure 3A, right panel) shows abasic sites in the RNA. Together, the results show that MPG binds to the R-loops upstream of *APOE* and forms RNA abasic sites.

**Figure 3.**
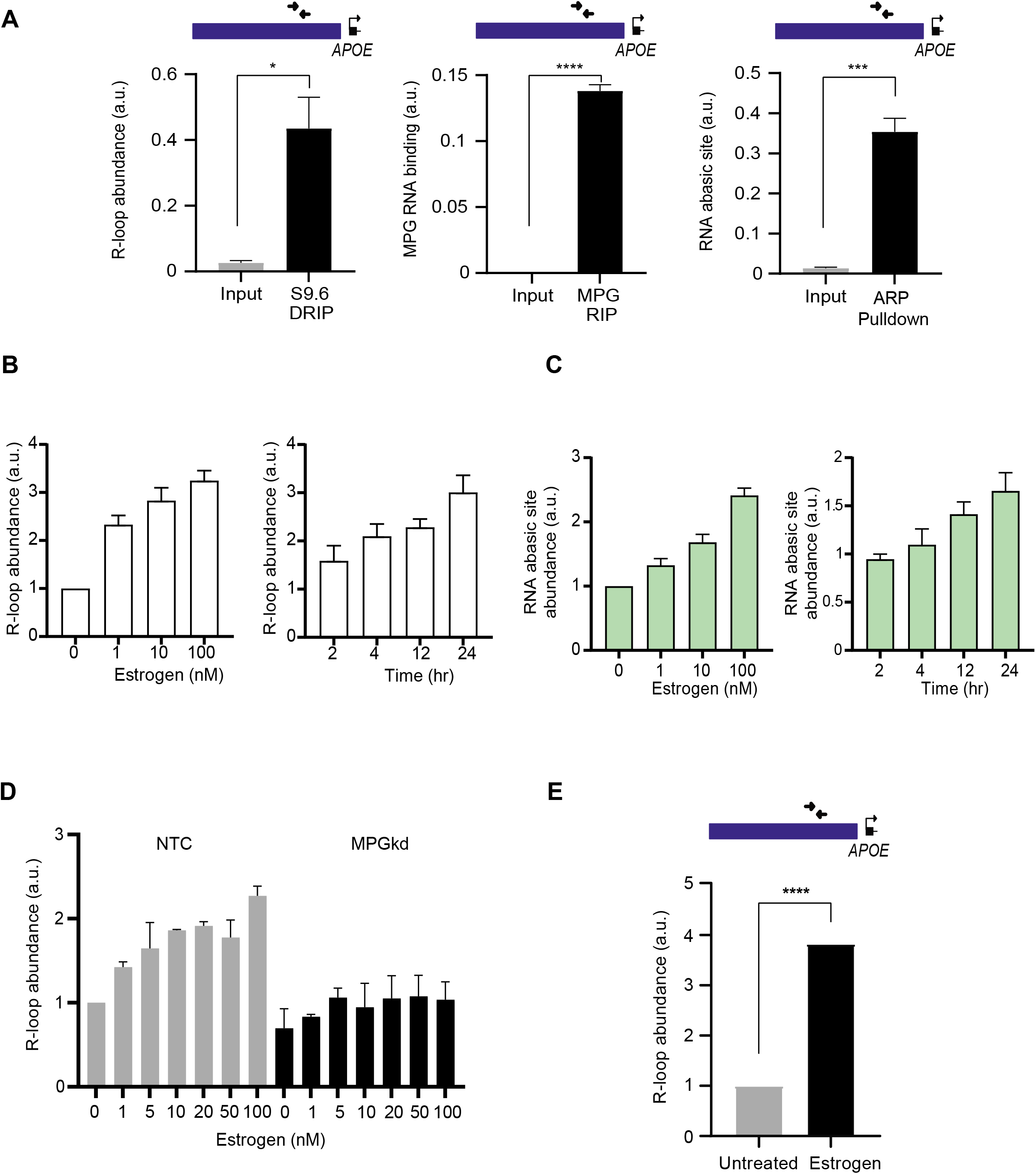
Upstream of *APOE*, RNA abasic sites forms in and stabilize the R-loops. (A) Data from S9.6 DRIP, MPG RNA-IP, and ARP RNA-pulldown followed by PCR are shown in arbitrary units (a.u.) Primers corresponding to the R-loops are shown in Figure 2A. S9.6 DRIP-PCR confirmed the location of the R-loops (N=3,*P<0.05, t-test, error bars=S.E.M.), MPG RIP-PCR shows binding of MPG to RNA of the R-loops (N=3, ****P<0.0001; t-test, error bars=S.E.M.), and ARP RNA-pulldown-PCR shows abasic sites in the RNA of the R-loops (N=3, ***P<0.001; t-test, error bars=S.E.M.). (B) R-loop abundance measured by S9.6 dot blot (see Methods) shows a dose-dependent increase of R-loops at 24 hours following treatment with estrogen (P<0.0001; ANOVA, error bars=S.E.M.) and time-dependent increase following 100 nM estrogen treatment (P<0.05; ANOVA, error bars=S.E.M.). (C) RNA abasic site abundance as measured by ARP-labeling shows a dose (N=3, ****P<0.0001; ANOVA, error bars=S.E.M.) and time (N=3, *P<0.05; ANOVA, error bars=S.E.M.) dependent increase following treatment with estrogen. Y-axis is fold-enrichment relative to control. (D) R-loop abundance was measured by S9.6 dot-blot in fibroblasts treated with scrambled siRNA (nontarget control, NTC) and siRNA against MPG, the cells were then given estrogen (0 to 100 nM) (N=2, error bars=S.E.M.). R-loops do not accumulate in response to estrogen when MPG is knocked down. (E) More R-loop are present upstream of *APOE* as measured by S9.6 DRIP-PCR in fibroblasts followed by estrogen treatments (100 mM). Location of the assessed R-loops is shown in the schematic and corresponds to the R-loops in Figure 2A (N=3; ****P<0.0001, t-test, error bars=S.E.M.).

To further examine the relationship between R-loops and RNA abasic sites, particularly whether there is a reciprocal relationship, we induced R-loops with estrogen (Stork et al., 2016). In primary fibroblasts, estrogen significantly (P<0.0001) increases R-loops (Figure 3B) and RNA abasic sites (Figure 3C) in a dose and time-dependent manner genome-wide. We then knocked down MPG and treated those fibroblasts with estrogen. Results show that when MPG is knocked down, R-loops do not accumulate in response to estrogen (Figure 3D), indicating that the stability of R-loops may depend on RNA abasic sites.

The estrogen treatment increases R-loops genome-wide, including upstream of *APOE* (Figure 3E). We assessed the effect of the stabilized R-loops on transcription upstream of *APOE* by performing PRO-seq in fibroblasts at different timepoints following estrogen treatment. The estrogen-induced R-loops led to increased pausing of RNA Pol II in the intergenic region, and we observed a time-dependent increase in RNA Pol II pausing (pausing index increased from 4.8 in resting cells to 11.8 after 6 hours in estrogen; Supplemental Figure 2).

Together, these results show that upstream of *APOE*, nascent RNA form R-loops, then MPG generates RNA abasic sites which likely stabilizes the R-loops that in turn pauses transcription of the intergenic RNA.

### m6A attracts MPG to R-loops

We next asked what attracts MPG to the RNA of the R-loops upstream of *APOE*. Given that MPG is a methylpurine glycosylase, we focused on methylated purines. According to Modomics (Boccaletto et al., 2018), there are 50 different types of methylated purines in RNA. We focused on m6A because it is highly abundant (Frye et al., 2016; Pan, 2013; Shi et al., 2019; Zaccara et al., 2019), and is found in R-loops (Abakir et al., 2020; Kang et al., 2021; Zhang et al., 2020).

We asked if m6As are present in the RNA in the R-loops upstream of *APOE* and if so, whether they are the target of MPG. We carried out RNA-IP with an antibody against m6A followed by PCR with the same primers used to assess MPG binding to the R-loops upstream of *APOE* in Figure 2A. The results show that m6As are found in the R-loops upstream of *APOE* (Figure 4A) and coincide with where MPG binds (Figure 3A, middle panel). The colocalization data suggest that m6As in the R-loops may be the substrate of MPG and therefore the precursor to RNA abasic sites.

**Figure 4.**
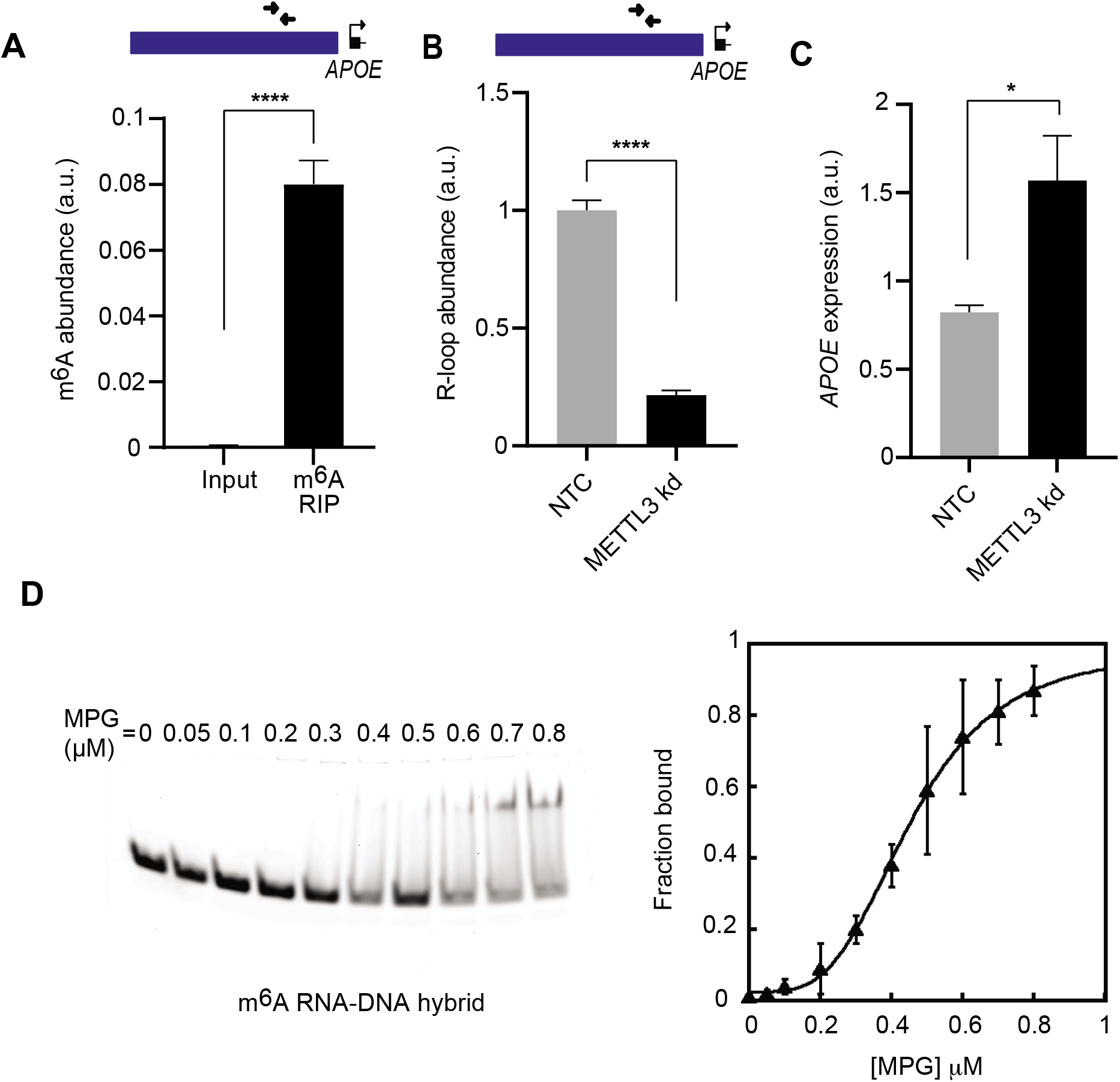
m6A is the precursor to RNA abasic site. (A) m6A in the R-loops upstream of *APOE* are identified by m6A-RIP followed by PCR with primers shown in the schematic and corresponds to the R-loops in Figure 2A (N=3, ***P<0.001; t-test, error bars=S.E.M.) m6A enrichment is shown in arbitrary units (a.u). (B) fewer R-loops upstream of *APOE* (N=3, ****P<0.0001; t-test, error bars=S.E.M.) and (C) higher *APOE* expression levels (N=3, *P<0.05; t-test, error bars=S.E.M.) in cells treated with siRNA targeting METTL3. Y-axis is fold-expression relative to NTC. (D) Representative blot showing gel shift from MPG binding to ^m6A^RNA-DNA hybrids (left). Binding data from gel shift experiments were fitted to the modified Hill equation (see Methods) with Kd=58 nM (right) (n=3, error bars=S.D.).

If m6A is a substrate of MPG, then the METTL3/14 complex that methylates adenosine to form m6A (Liu et al., 2014; Wang et al., 2016) should also regulate *APOE* expression. We knocked down METTL3 (Supplementary Figure 3A) and measured the abundance of the R-loop upstream of *APOE* and *APOE* expression. In cells whose METTL3 was knocked down, we found significantly (P<0.001) fewer R-loops (Figure 4B) and significantly (P<0.05) higher *APOE* expression (Figure 4C) without affecting the expression of MPG (Supplemental Figure 3B). Hence, METTL3 knockdown phenocopies MPG knockdown, and likely acts upstream of MPG in the regulation of *APOE*.

To determine if m6A is indeed a substrate of MPG, we carried out assays to assess if MPG can incise m6A from RNA. Electromobility shift assays show that MPG had a strong affinity (Kd=58 nM) for the hybrid with m6A RNA (Figure 4D). Next, we incubated the RNA-DNA hybrids with MPG followed by AP endonuclease 1 (Liu et al., 2020) which cleaves the sugar-phosphate backbone at the abasic site. The results showed that MPG incised m6A when the RNA strand of an RNA-DNA hybrid contains an m6A, but not when the m6A is in the DNA strand of the RNA-DNA hybrid (Supplementary Figure 3C). Thus, m6A is removed by MPG to form an RNA abasic site. Taken together, m6As are the precursor of RNA abasic sites in the R-loops upstream of *APOE*.

### A non-coding RNA upstream of *APOE*

Thus far, we have discussed the R-loops that form upstream of *APOE*, without characterizing the transcription and function of the RNA. Our PRO-seq data from primary fibroblasts show that there is more active transcription in the intergenic region between *TOMM40* and *APOE*. There are more RNA Pol II and therefore higher transcription in the intergenic region than in *TOMM40* and *APOE;* thus, the RNA is likely an independent transcript and not part of *TOMM40* or *APOE* (Supplementary Figure 4A). With data from CoPRO (Figure 5A) (Tome et al., 2018) and PRO-cap (Supplementary Figure 4B) (Core et al., 2014; Mahat et al., 2016) of the intergenic region, we identified capped RNA thus the 5’ end of the RNA. The capped RNA coincides with a region with active transcription as shown by PRO-seq (Figure 5A, Supplementary Figure 4B), confirming that the transcript identified in CoPRO and PRO-cap is the same as that identified by PRO-seq. We then searched for a polyadenylation motif and did not find one in this RNA.

**Figure 5.**
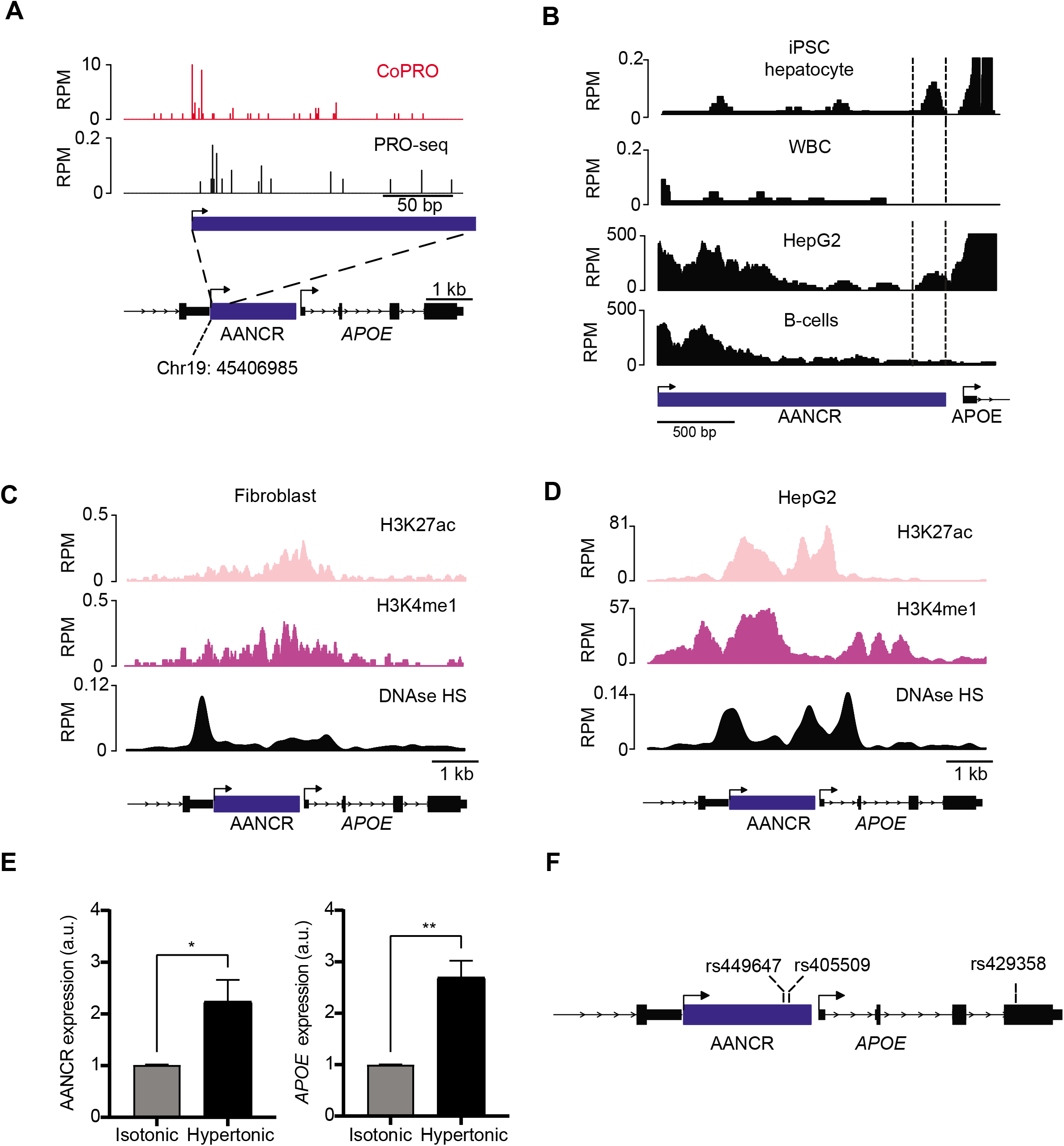
AANCR functions as an R-loop-dependent enhancer. (A) CoPRO identified capped RNA in the intergenic region upstream of *APOE*, the cap coincides with nascent transcription from PRO-seq. The transcription start site of the non-coding RNA, AANCR, is marked based on the cap location. Genomic coordinates (hg19) are indicated. (B) RNA-seq data showing full-length AANCR in iPSC-derived hepatocytes and a partial transcript of AANCR in white blood cells (WBC). BRU-seq data show fulllength AANCR in HepG2 liver cells and partial AANCR transcript in cultured B-cells. (C,D) At the AANCR locus, ChIP-seq results show enhancer marks H3K27ac and H3K4me1, and DNase mapping results show DNase hypersensitivity in fibroblasts (C) and HepG2 cells (D). (E) Expression of full-length AANCR (N=3, P<0.05:t-test, error bars=S.E.M.) and *APOE* (N=3,**P<0.01;t-test, error bars=S.E.M.) are induced in response to hypertonic stress with the addition of 50 mM NaCl to the media culture of HK-2 renal proximal tubule cells. (F) Schematic of SNPs in AANCR and *APOE*.

Next, we asked if this capped RNA is noncoding. Sequence analysis by BLASTX (Gish and States, 1993) and PFAM (Bateman et al., 2004) did not identify similar proteins. Additionally, we analyzed the sequence by several algorithms that test for coding potentials (CPAT, CPC, CNIT(Guo et al., 2019; Kong et al., 2007; Wang et al., 2013)). These analyses determined that this is a noncoding RNA (Table 1). Together, the results show that upstream of *APOE* is a capped noncoding RNA, which we have named APOE associated noncoding RNA (AANCR).

**Table 1.** The RNA upstream of *APOE* is a noncoding RNA.

| Method | Results |
| --- | --- |
| BLASTX | Only 3 sequences showed >80% identity, two are hypothetical proteins, one is a low-quality protein. |
| PFAM | No significant hits by PFAM-A HMM and GA cutoffs |
| CPAT | Fickett score(Fickett, 1982)=0, hexamer score=0, coding potential=0.003; classified as noncoding |
| CNIT | Score=0.31, classified as noncoding |
| CPC | Coding probability=0.04, classified as noncoding |

Our data show that in fibroblasts, R-loops pause RNA Pol II elongation so AANCR is only partially transcribed, and *APOE* is not expressed or expressed at a very low level (Figure 2A and Supplemental Figure 2). Next, we extended the analysis by performing RNA-seq of other cell types, including liver cells that express *APOE* and white blood cells that do not express *APOE*, to assess if the length of AANCR correlates with *APOE* expression. Figure 5B shows in iPSC-derived hepatocytes (Carpentier et al., 2014), AANCR is transcribed as a full-length transcript, but in white blood cells (from the same individual from which the hepatocytes were derived), AANCR is only partially transcribed. To confirm that in cells with full-length AANCR, *APOE* is expressed but in cells with partial transcription of AANCR, *APOE* is not expressed, we turned to BRU-seq data (Paulsen et al., 2013) in the ENCODE data portal (Davis et al., 2018). BRU-seq data also show full-length AANCR in liver cells that express *APOE*, but the partially transcribed AANCR is found in cultured B-cells where *APOE* is not expressed (Figure 5B). In the liver cells where AANCR is full-length, we did not find R-loops, m6As, or RNA abasic sites (Supplemental Figure 4C) confirming that without stable R-loops, AANCR is transcribed as a full-length RNA that enhances *APOE* expression.

In summary, we have discovered a noncoding RNA, AANCR, upstream of *APOE* and that AANCR is regulated by R-loops with RNA base modifications. In liver cells, AANCR is full-length and *APOE* is expressed; in contrast, in fibroblasts and B-cells, R-loops pause RNA Pol II transcription so AANCR is only partially transcribed, and *APOE* is not expressed.

### AANCR is an enhancer RNA

Next, we assessed if this noncoding RNA is an enhancer of *APOE*. The ENCODE registry of candidate cis-regulatory elements (Moore et al., 2020) had identified the region upstream of *APOE* as a potential enhancer based on DNase hypersensitivity and histone modification. GeneHancer also annotated AANCR as an elite enhancer based on the criteria that the region has features of enhancer chromatin and a strong enhancer-gene association (Fishilevich et al., 2017). In primary fibroblasts, we performed ChIP-seq to assess if the AANCR region has features consistent with an enhancer. In the AANCR region, we found H3K27ac and H3K4me1 marks, and DNase hypersensitive chromatin that characterize enhancers (Heintzman et al., 2007, 2009) (Figure 5C). In HepG2 cells where *APOE* is expressed, as expected this region is marked with open chromatin and enhancer marks, H3K27ac and H3K4me1 (Figure 5D). Thus, AANCR is an enhancer RNA, and the region that compasses AANCR and *APOE* is poised for transcription even in cells where *APOE* is not expressed.

Next, we asked whether AANCR physically interacts with *APOE*. Given that AANCR and *APOE* are located adjacent to each other, we turned to the technique of Micro-C (Krietenstein et al., 2020) that maps chromatin folding at high resolution. The results showed that AANCR is in a ~18 kb topology-associated domain with *APOE* and the downstream gene *APOC1* (Supplemental Figure 4D). In contrast, *TOMM40* which is 12 kb upstream of AANCR is outside of that topology-associated domain. Following MPG knockdown, like *APOE, APOC1* is significantly induced (P<<0.001, 17-fold), whereas *TOMM40* is only slightly increased (1.4-fold) (Supplemental Figure 4E). Using the CoXpresDB (Obayashi et al., 2019) which allows users to query co-expression in more than 25,000 datasets from microarray and RNA-seq experiments in the public data bank (Kodama et al., 2018), we found that the expression levels of *APOE* and *APOC1* are highly correlated, while *APOE* and *TOMM40* expression are much less correlated (Supplementary Figure 4F). Thus, our finding that AANCR is an enhancer of *APOE* and *APOC1* is a general phenomenon, beyond the cell types that we examined in this study.

Together, we have identified a non-coding RNA, AANCR, which enhances *APOE* and *APOC1* expression. In cells where base modifications stabilize R-loops, AANCR is only partially transcribed and cannot promote *APOE and APOC1* expression, while in cells such as those in the liver, AANCR is a fulllength enhancer RNA that induces transcription and expression of *APOE* and *APOC1*.

### AANCR and *APOE* expression is stress-responsive

Data such as DNase hypersensitivity from the ENCODE consortium and in our labs show that the *APOE* region is poised for expression. This is somewhat surprising since *APOE* is known to be expressed only in several cell types, particularly liver and macrophages. We posit that *APOE* is poised to be transcribed in response to stress. We treated renal proximal tubule cells with hypertonic salt and measured AANCR and *APOE* expression. Under osmotic stress, renal proximal tubule cells express full-length AANCR and consequently induce *APOE* expression (Figure 5E). Thus, AANCR acts as an enhancer to facilitate *APOE* expression in response to cellular stress.

### Sequence variants in AANCR influence *APOE* expression and susceptibility to Alzheimer’s Disease

Since AANCR regulates *APOE* expression and the ε4 allelic form of *APOE* is a major risk factor for Alzheimer’s Disease (Corder et al., 1993), we wondered if AANCR may affect susceptibility to Alzheimer’s Disease. As a complex disease, there are multiple contributing factors, besides the ε4 variant, other genetic factors have been identified, including those upstream of *APOE* where AANCR is encoded (Figure 5F). Studies have reported evidence for population association of rs449647 (also referred to as −491) with Alzheimer’s Disease (Bullido et al., 1998; Lambert et al., 1998; Wang et al., 2000). The GWAS catalog also shows multiple studies that have identified significant allelic association of SNPs in the AANCR region including rs449647 and rs405509 with Alzheimer’s Disease (Table S1A). A study of 42,034 patients and 272,244 controls in the United Kingdom found significant association (P <10^-50^) of these SNPs with Alzheimer’s disease (Marioni et al., 2018). To assess how the sequence variants in AANCR affect susceptibility to dementia, we asked if the genetic variants affect the function of AANCR and therefore *APOE* expression. Using data from the GTEx consortium (Ardlie et al., 2015), despite the small sample size of a few hundred samples, we found that SNPs in the AANCR region are significantly associated with *APOE* expression in the liver and the brain, including the hippocampus (Table S1B). While *APOE* expression is important in understanding transcriptional regulation, it is not practical to measure *APOE* gene or protein expression in tissues from many individuals, so we asked if our finding can be extended to plasma APOE level, a more feasible clinical measurement. We turned to a Danish study that has measured plasma APOE of 106,652 individuals (Rasmussen et al., 2018). Their results show evidence for (P<10^-6^) association of SNP variants, including rs449647 in AANCR with plasma APOE level. Together, the genetic data confirm the role of AANCR in regulating the transcription of *APOE*. The genetic variants in AANCR that contribute to individual differences in *APOE* expression also affect risk of developing Alzheimer’s Disease.

Next, we assessed if polymorphisms in AANCR contribute additional risk factors to Alzheimer’s disease beyond the ε4 variant in *APOE*. Despite the proximity of AANCR and *APOE*, the extent of linkage disequilibrium is modest. From the different populations studied by the HapMap Consortium (International HapMap Consortium et al., 2007), the r^2^ of rs449647 (in ANNCR) and rs429358 (in *APOE*) are 0.054 in Western Europeans, 0.068 in Yoruba people from Idaban, 0.10 in Japanese from Tokyo and 0.08 in Han Chinese from Beijing. Thus, the allelic association of the SNPs in the AANCR regions with *APOE* expression and Alzheimer’s disease are most likely independent of those of *APOE*4. This suggests that by regulating *APOE* expression, AANCR is a modifier of Alzheimer’s disease.

## Discussion

Although there have been tremendous advances in our understanding of RNA, we are still a long way from knowing its components and likely most of its function. RNA was once considered simply as an essential intermediary between DNA instructions and protein synthesis. We now know that RNA is much more than a go-between DNA and protein, and that RNA carries important regulatory information. RNAs are also more varied than originally thought. It is now known that nearly every nucleotide in DNA is transcribed into RNA (ENCODE Project Consortium et al., 2007; Hangauer et al., 2013), so in addition to mRNA and tRNA, there are myriad noncoding RNAs (Cech and Steitz, 2014; Mattick, 2018; Mercer et al., 2009). Many noncoding RNAs are found in intergenic regions where SNP alleles are associated with diseases thus identifying these RNAs and understanding their regulatory functions are critically important (Tak and Farnham, 2015; Zhang and Lupski, 2015). Co-transcriptionally and post-transcriptionally, the bases and sugar of RNA are modified, transcripts are spliced and polyadenylated with different lengths of adenosine. These processing steps add to the complexity of RNA by generating different transcripts from the same DNA template. The processing of RNA differs by cell-type and cellular environments resulting in a vast array of RNA species with different regulatory roles.

This study was motivated by our previous work that uncovered abasic sites in RNA that are present in R-loops. The discovery of DNA abasic sites and the enzymes such as methylpurine glycosylase which form and process abasic sites, led to the elucidation of base excision repair in response to DNA damage (Lindahl, 1993). However, the abasic sites in RNA differ from those in DNA due to the additional hydroxyl group. Studies have shown that RNA with abasic sites is more stable than DNA with abasic sites, as cleavage at the abasic site in RNA is at least 17 times less likely than at the abasic site in DNA (Küpfer and Leumann, 2007). RNA abasic sites are not rare, by mass spectrometry, we found that there are about 4 RNA abasic sites per million ribonucleotides (Liu et al., 2020); so with over 50 billion ribonucleotides in a cell, there are hundreds of thousands of RNA abasic sites in each cell. Given that RNA with abasic sites is stable, and there are many abasic sites in RNA, it seems critical to determine their regulatory roles. Additionally, tens of thousands of R-loops have been mapped genome-wide (Crossley et al., 2020; Ginno et al., 2012; Sanz et al., 2016; Wahba et al., 2016). We are just beginning to understand some of their features and functions with many aspects of R-loops remaining to be explored, such as the stability of R-loops, and the relationship between stability and function (Belotserkovskii and Hanawalt, 2022; Castillo-Guzman and Chédin, 2021).

Here, we examined a group of R-loops with RNA abasic sites and uncovered that their stability allows them to pause RNA Polymerase II transcription. We identified a noncoding RNA, AANCR, and show that its transcription and function are dynamically regulated by RNA sequence modifications and the nucleic acid structure, R-loop. In the R-loops of the noncoding RNA, AANCR, the RNA is modified by N6-adenine methylation and then the resultant N6-methyladenines are removed by methylpurine glycosylase to become abasic sites which stabilize the R-loops. In some cell types, the stabilized R-loops pause RNA Pol II and prevent the synthesis of the full-length enhancer RNA, yet upon stress, the R-loops resolve, the enhancer RNA is fully transcribed to allow rapid activation of *APOE*. In other cell types and cellular conditions, the R-loops are not modified and do not persist, thus the noncoding RNA is transcribed into a full-length enhancer that activates *APOE* expression. The transcribing RNA Pol II synthesizing AANCR is paused by the modified R-loop without the NELF protein complex that pauses transcription in the proximal promoter regions. This nucleic-acid-mediated pausing keeps AANCR poised for rapid transcriptional responses. The R-loops can resolve quickly to allow RNA Pol II to complete the synthesis of AANCR and activate its target genes. Like the hammerhead ribozyme whose catalytic function is dependent on its structure (Pley et al., 1994; Scott et al., 1995), AANCR is also dependent on its structure to function as an enhancer of *APOE* and *APOC1* expression. In fibroblasts at basal state, there are over 6,000 R-loops where MPG binds thus nucleic-acid-mediated transcriptional pausing of AANCR is likely a general mechanism that regulates enhancer RNAs.

In studying the RNA abasic sites and R-loops in AANCR, we revealed the mechanism that activates *APOE* transcription and expression. Previously, the induction of *APOE* in response to lipid load is well characterized, it involves the 3’ enhancer elements (Shih et al., 2000) and LXR/RXR signaling (Laffitte et al., 2001). Here, we show how cell-type and stress-responsive expression of *APOE* are regulated. Understanding the regulation of *APOE* expression is critical given its role in lipid transport (Mahley, 1988), maintenance of cognitive function (Corder et al., 1993; Mahley and Huang, 2012; Roses, 1996), and response to infection (Jiang and Luo, 2009; Kuo et al., 2020) as well as immunotherapies (Pencheva et al., 2014; Tavazoie et al., 2018). Our study shows that sequence variants in AANCR explain individual variation in *APOE* expression and APOE plasma level. The variants in AANCR region that affect APOE level also confer susceptibility to Alzheimer’s disease. Based on the extent of linkage disequilibrium between the variants in AANCR and *APOE*, the effect of polymorphisms in AANCR is independent of the risk conferred by the *APOE* ε4 allele. By regulating *APOE* expression, AANCR acts as a modifier of Alzheimer’s disease.

Genetic studies of Alzheimer’s disease found that the A-allele of −491 SNP (rs449647) is the risk allele (Bullido et al., 1998; Lambert et al., 2013) and in the GTEx data, the A-allele is associated with lower *APOE* expression in different cell types including the hippocampus. While some studies have shown that Alzheimer’s patients have low plasma APOE levels (Bertrand et al., 1995; Rasmussen et al., 2018), other studies report more complex findings including higher APOE levels in the patients (Koch et al., 2020; Laws et al., 2002; Lehtimäki et al., 1995; Taddei et al., 1997). One would expect that the protective effect of APOE levels may differ depending on the *APOE* genotype, where higher APOE3 may be protective but higher APOE4 level is not. Future studies can assess how variation in AANCR and *APOE* interact to affect Alzheimer’s disease and those results may guide the modulation of AANCR in RNA-based therapeutics in the prevention and treatment of Alzheimer’s disease.

In conclusion, we identified a regulatory mechanism where the METTL3/METTL14 protein complex coalesces with methylpurine glycosylase forming N6-methyladenines which are then cleaved resulting in RNA abasic sites on R-loops to pause transcription. We illustrated how this pathway that includes RNA base modifications and R-loops regulates the expression of *APOE*. We surmise that RNA modification and RNA structure play a critical role in the dynamic regulation of gene expression. Studies of nucleic acid sequence and structure will advance the basic understanding of gene regulation and the genetic basis of complex diseases.

---

**Supplemental Figure 1.**
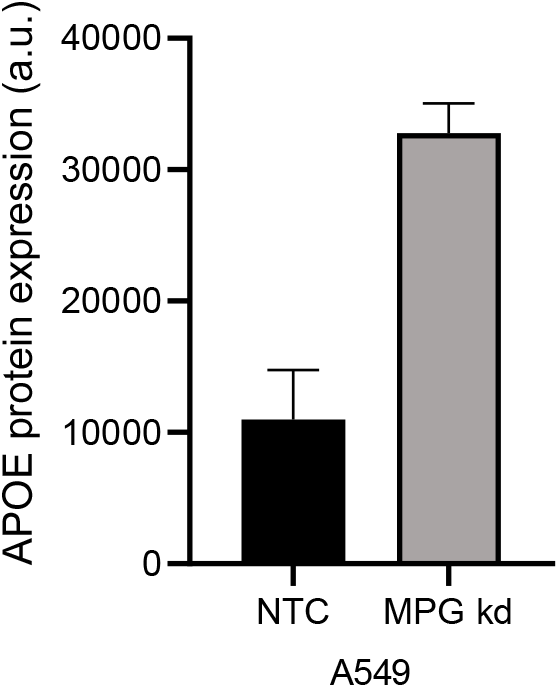
MPG knockdown induces *APOE* expression in A549, lung epithelial cells. *APOE* expression levels as measured by RT-PCR in A549 with and without MPG knockdown (day 7) are plotted (N=2, *P<0.05, t-test, error bars=S.E.M.).

**Supplemental Figure 2.**
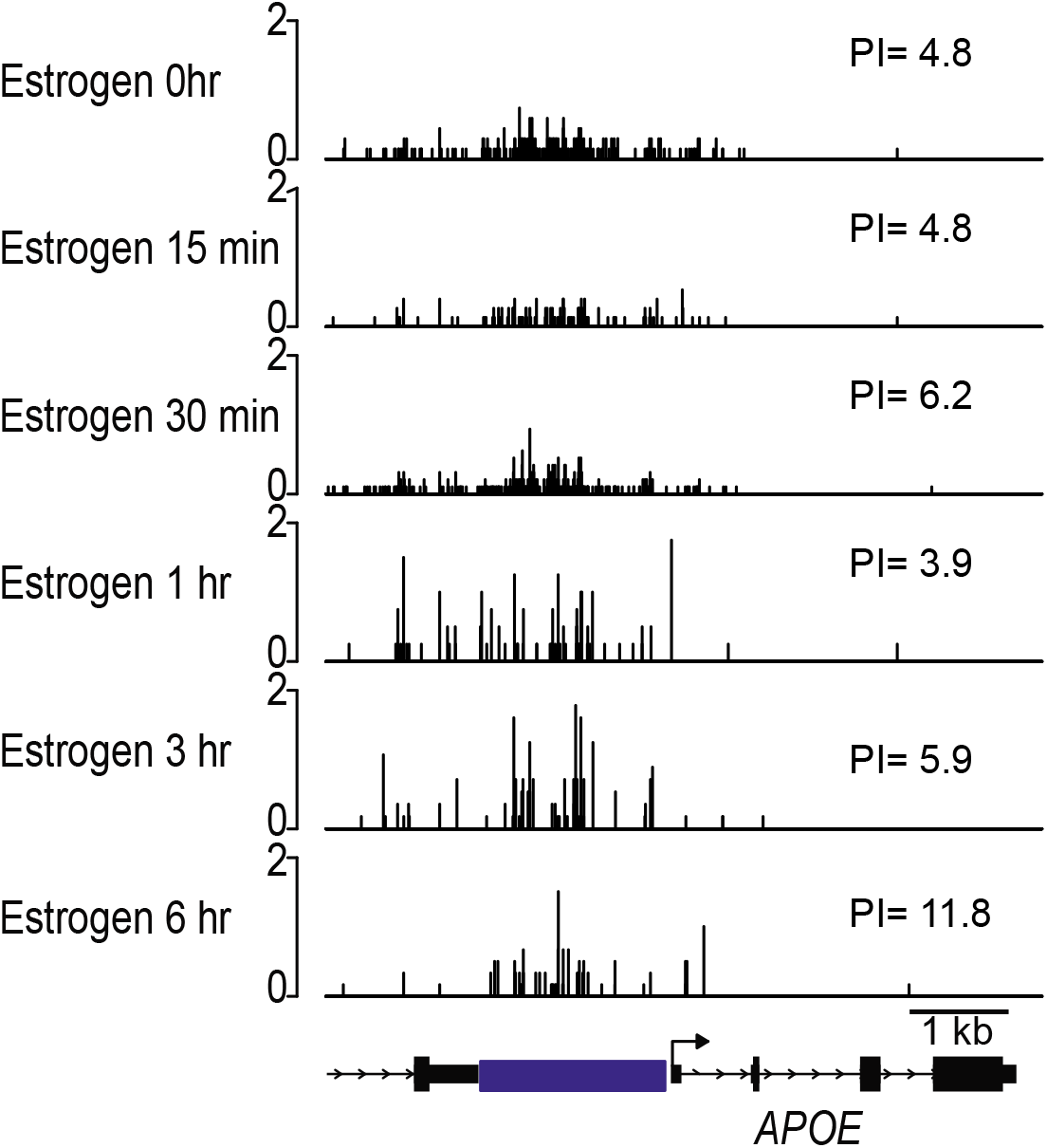
Estrogen increases RNA Pol II pausing upstream of *APOE*. PRO-seq carried out using fibroblasts treated with estrogen (100nM) for different periods of time, data shown for the intergenic region upstream of *APOE*. Pausing index (PI) calculation (see Methods) shows that estrogen increases pausing of RNA Pol II.

**Supplemental Figure 3.**
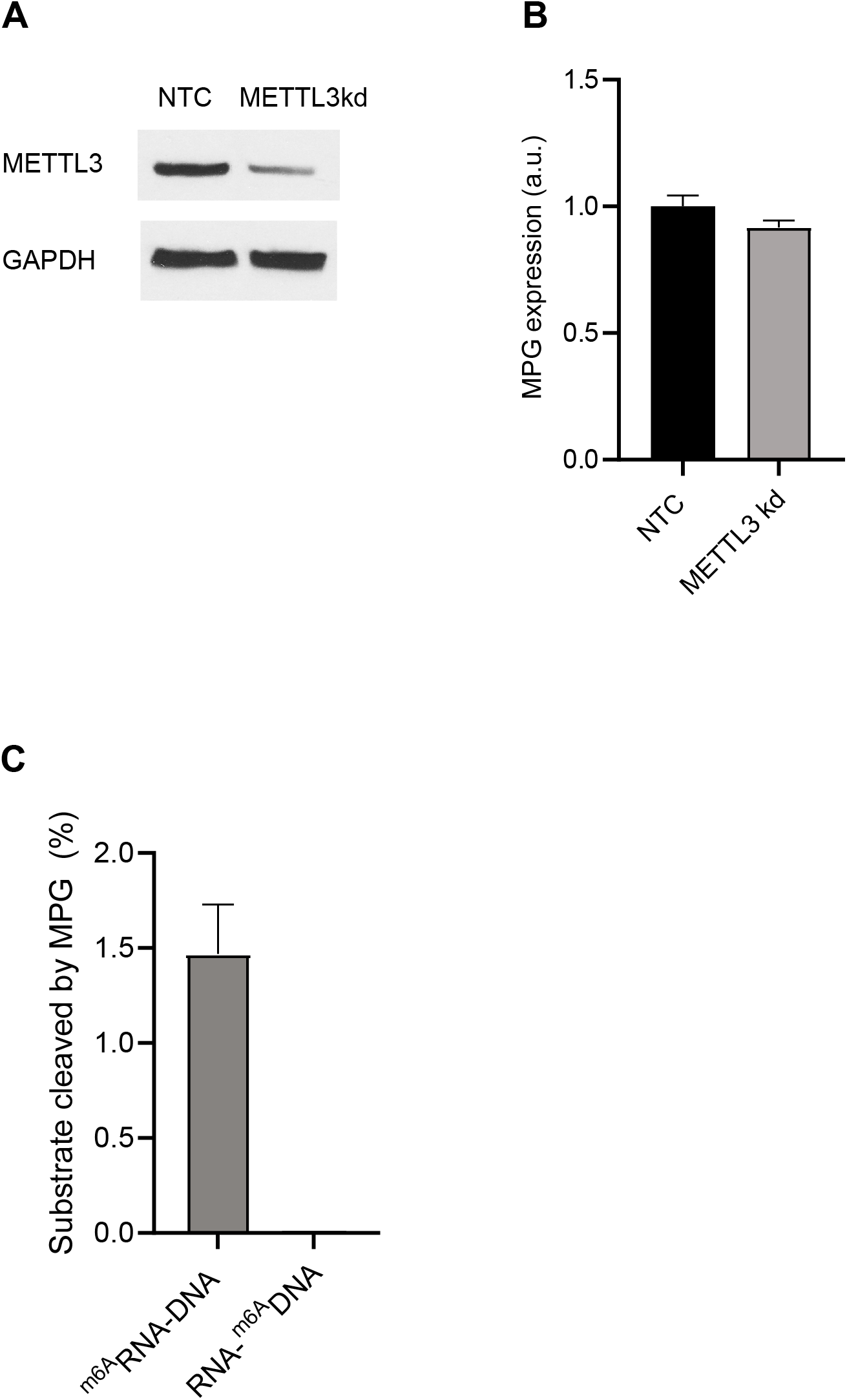
m6A is a precursor of the RNA abasic site upstream of APOE. (A) Representative immunoblot with anti-METTL3 antibody and GAPDH as loading control confirm that METTL3 expression is reduced following siRNA knockdown. (N>3) (B) MPG expression in fibroblast with and without METTL3 siRNA knockdown, as measured by RT-PCR. (N=2, P>0.5, error bars=S.E.M.) (C) Percentage of cleaved ^m6A^RNA-DNA or RNA-^m6A^DNA substrates by MPG, as assessed by addition of MPG to the substrates followed by addition of APE1 (N=3, error bars=S.E.M.).

**Supplemental Figure 4.**
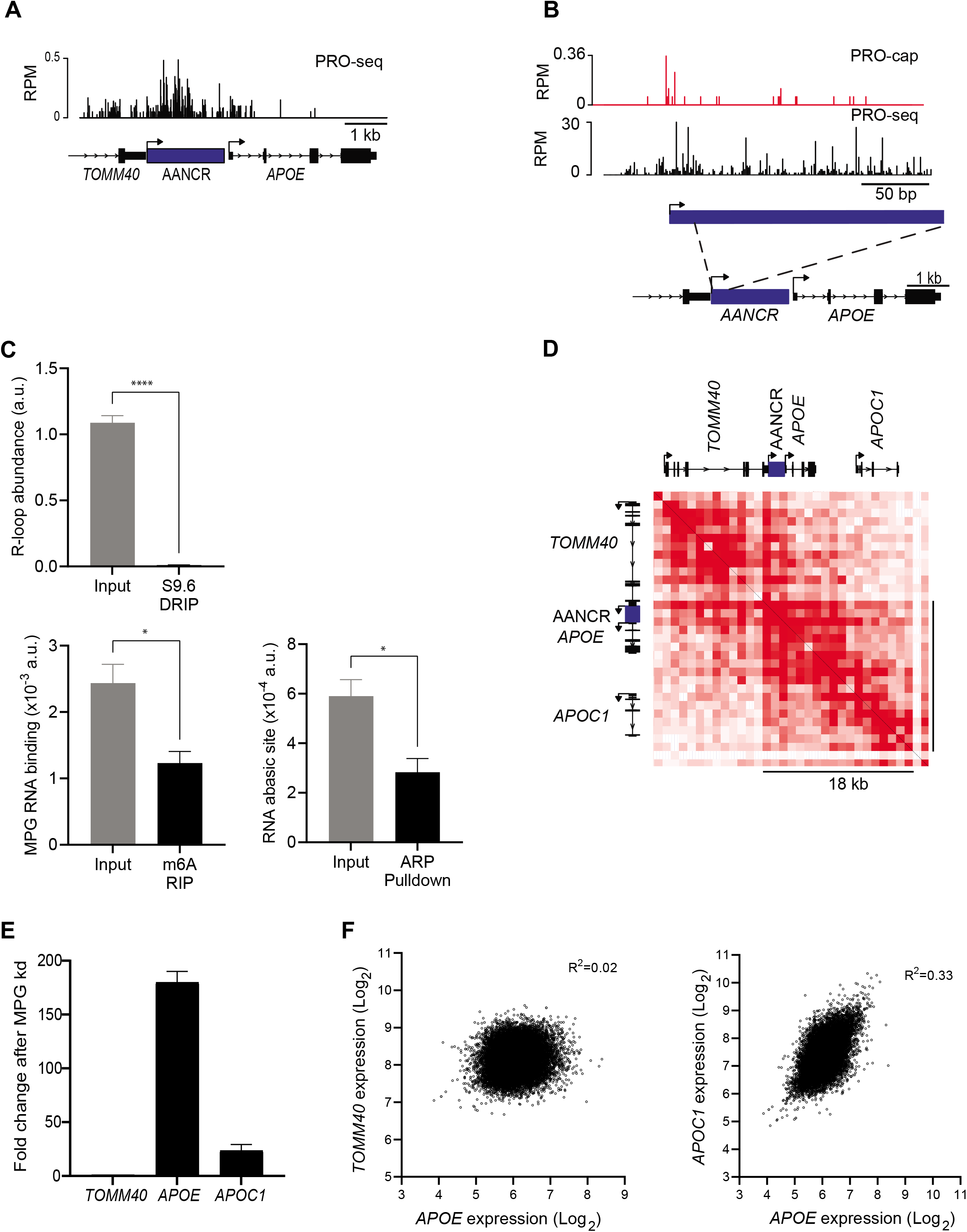
AANCR, *APOE* and *APOC1* are in the same topological domain. (A) PRO-seq from fibroblasts showing more active transcription at AANCR as compared to *TOMM40* or *APOE*. Y-axis is RPM. (B) PRO-cap (upper panel) and PRO-seq (lower panel) from K562 lymphoblasts identify a capped transcript in the intergenic region upstream of *APOE*. Y-axis is RPM. (C) Data from S9.6 DRIP, MPG RNA-IP and ARP RNA-pulldown followed by PCR in HepG2 are shown in arbitrary units (a.u.). Primers corresponding to the R-loops are shown in Figure 2A. MPG RIP-PCR and ARP RNA-pulldown-PCR did not identify m6A or abasic sites in the RNA of the R-loops (N=3, error bars=S.E.M.). (D) Hi-C contact map of the *TOMM40-APOC1* locus reveals AANCR, *APOE*, and *APOC1* are within an 18 kb loop topologically-associated domain. The *in situ* Hi-C chromatin structure from transformed foreskin fibroblasts at 1kb resolution were downloaded from the UCSC genome browser (chr19:44,889,969-44,922,968, hg38). (D) Expression of *TOMM40, APOE*, and *APOC1* as measured by RNAseq before and after MPG knockdown (day 7) in fibroblasts show induction of *APOE* and *APOC1* that are in the same chromatin loop with AANCR, but not *TOMM40* (N=2, error bars=S.E.M.) which is in a different topological domain. (E) Comparison of gene expression between *APOE* and *TOMM40* (left), or *APOE* and *APOC1* (right) indicate *APOE* and *APOC1* are correlated across datasets (N=25,362) in COXPRESdb version 8.0. X and Y axis are relative expression levels (Log2).

**Table S1A.** Examples of SNPs in AANCR region that show allelic association with Alzheimer’s Disease.

| SNP in AANCR | P-value* (Alzheimer's Disease) | Study |
| --- | --- | --- |
| rs449647 | $2 \times 10^{-58}$ | 42,034 British ancestry individuals with parental history of Alzheimer's disease, at least 272,244 individuals with no parental history of Alzheimer's disease (Marioni et al., 2018) |
| | $8.7 \times 10^{-18}$ | Transethnic LAOD (Jun et al., 2017) |
| | $2.6 \times 10^{-31}$ | Meta-analysis of 74,046 individuals (Lambert et al., 2013) |
| rs405509 | $1 \times 10^{-8}$ (males)<br>$4 \times 10^{-14}$ (females) | 952 European ancestry male cases, 1,789 European ancestry female cases, 6,337 European ancestry male controls, 8,402 European ancestry female controls (Nazarian et al., 2019) |
| | $3 \times 10^{-44}$ | ADGC GWAS Stage 1 (Naj et al., 2011) |
| | $6.1 \times 10^{-12}$ | ADGC African American (Reitz et al., 2013) |

\*Obtained from GWAS catalog and the National Institute of Aging Genomics of Alzheimer's Disease database.

**Table S1B.**
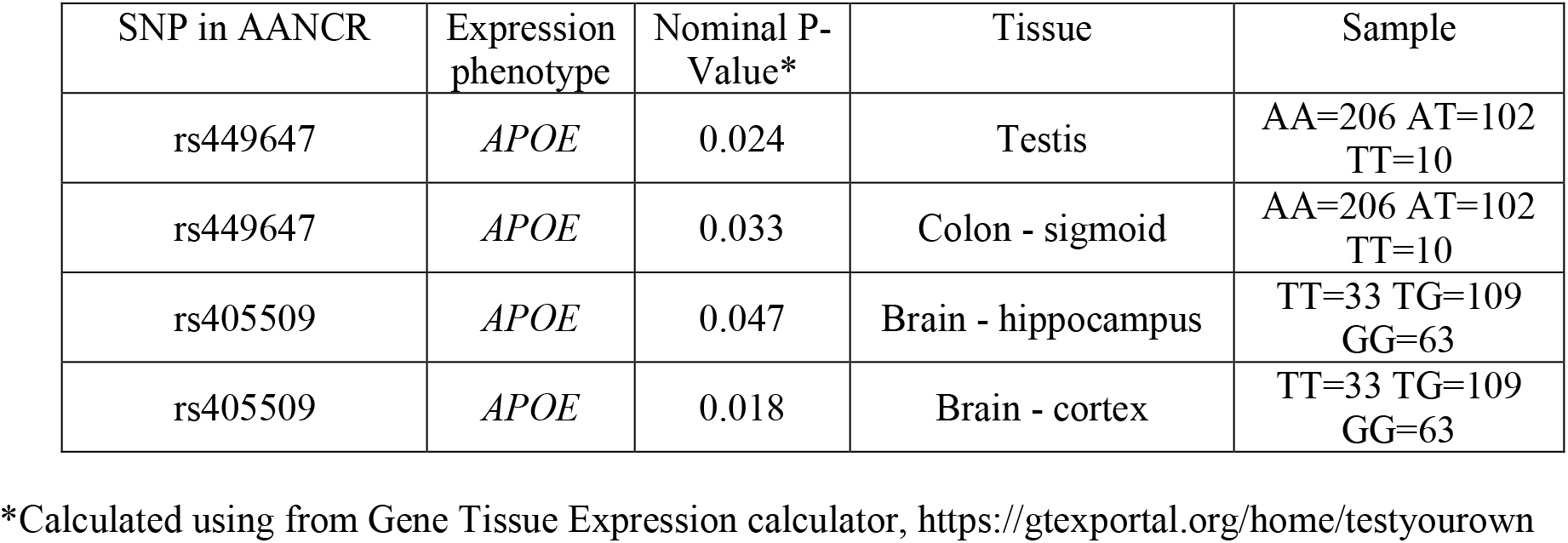
Association results for SNPs in AANCR with *APOE* expression level as phenotype.

| SNP in AANCR | Expression phenotype | Nominal P-Value* | Tissue | Sample |
| --- | --- | --- | --- | --- |
| rs449647 | *APOE* | 0.024 | Testis | AA=206 AT=102<br>TT=10 |
| rs449647 | *APOE* | 0.033 | Colon - sigmoid | AA=206 AT=102<br>TT=10 |
| rs405509 | *APOE* | 0.047 | Brain - hippocampus | TT=33 TG=109<br>GG=63 |
| rs405509 | *APOE* | 0.018 | Brain - cortex | TT=33 TG=109<br>GG=63 |

\*Calculated using from Gene Tissue Expression calculator, https://gtexportal.org/home/testyourown

**Table S2.**
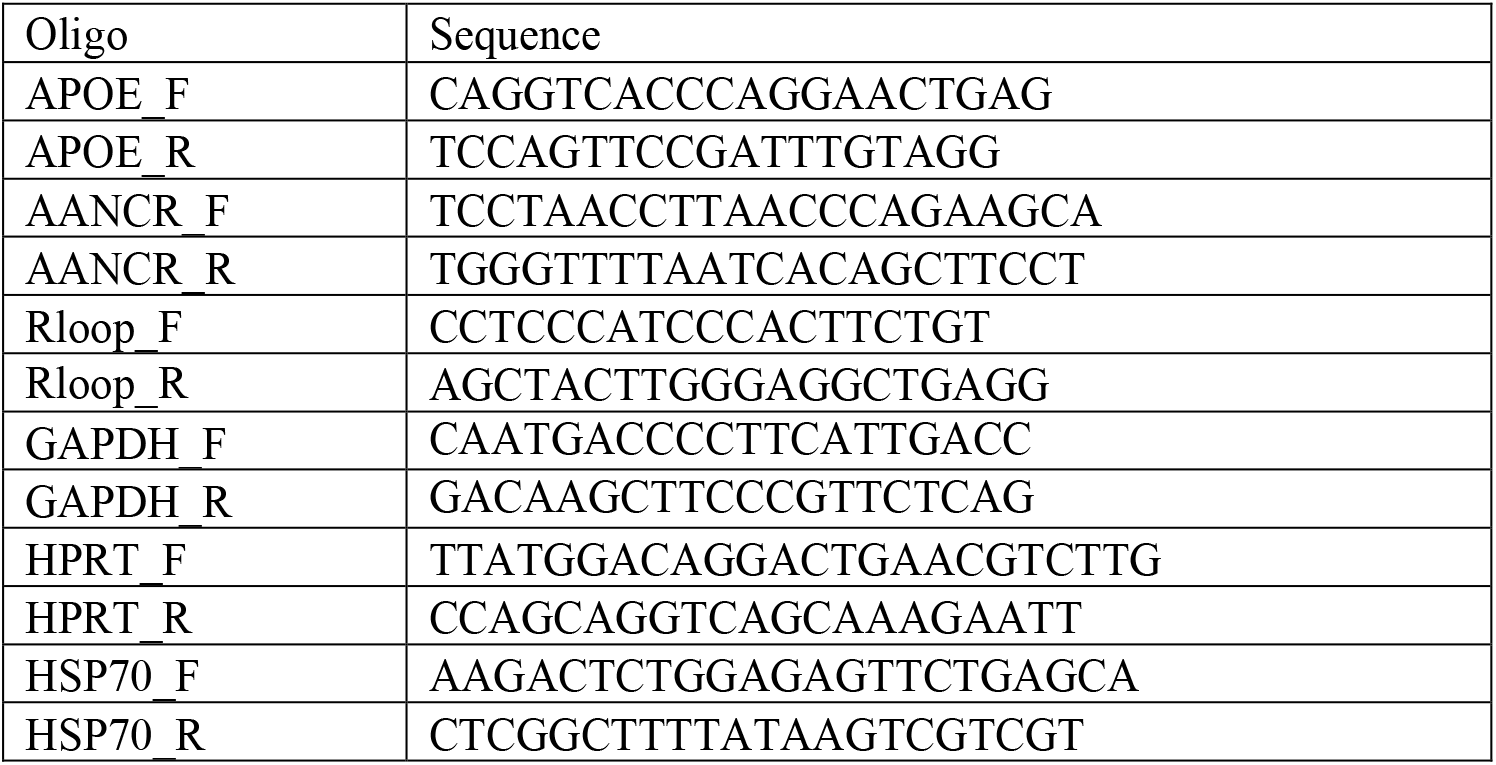
Oligonucleotides for qPCR

**Table S3.**
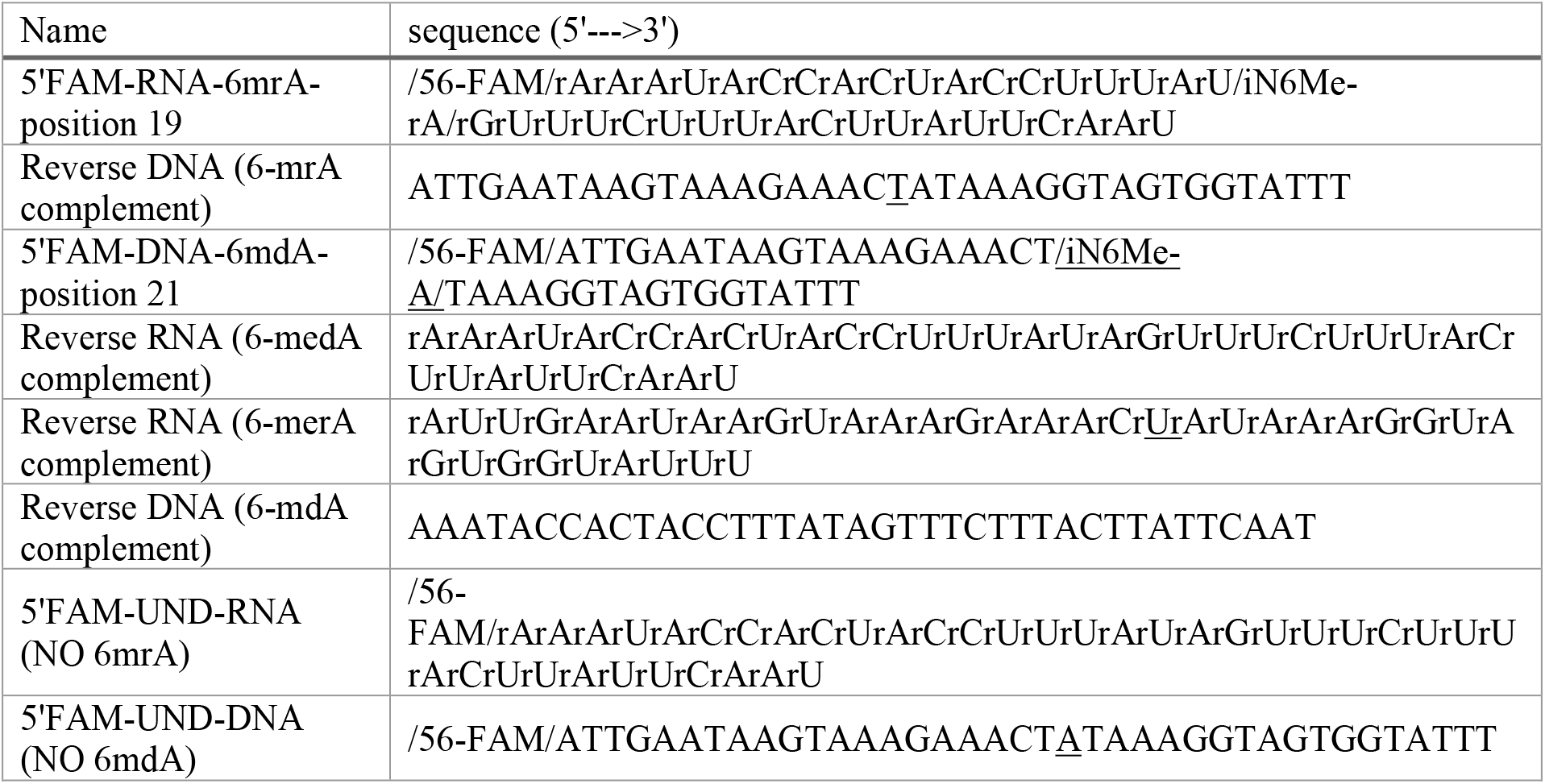
Substrates used for MPG assay.

## Acknowledgments

We thank Annelise Comai for assistance with data processing and analysis. We thank Drs. William Copeland, Paul Doetsch, Philip Hanawalt for comments and suggestions.

## Data and material availability

The deep sequencing data reported in this paper have been deposited in the NCBI sequence read archive, PRJNA801792 and the NCBI database of Genotypes and Phenotypes archive, Phs001322.v2.p1. Downloaded data are listed in the Key Resources Table.

## Methods

### Cell culture

Foreskin fibroblasts from healthy newborns were cultured in MEM medium (Thermo-Fisher) supplemented with 10% fetal bovine serum, 1% L-glutamine, and 1% penicillin-streptomycin. HK-2 cells (ATCC) were cultured in DMEM/F12 supplemented with 10% fetal bovine serum and 1% penicillin-streptomycin. A549 (ATCC) and HepG2 (ATCC) cells were cultured in DMEM supplemented with 10% fetal bovine serum and 1% penicillin-streptomycin. All cells were grown at 37°C with 5% CO2 and were passaged every 72 hours using Trypsin-EDTA (0.05%). Where indicated media were supplemented with estrogen (dissolved in ethanol) and equal volume of vehicle. For hypertonic stress experiment, media was supplemented with 50 mM NaCl and added to confluent HK-2 cells for 6 hours before RNA isolation.

### Chromatin Immunoprecipitation

Foreskin fibroblasts were cross-linked with 1% formaldehyde for 10 min. Cross-linking was stopped with 2.5M glycine for 5 min. Nuclei were isolated by rotating crosslinked cells for 10 min at 4°C in 5 ml lysis buffer 1 (50 mM Hepes pH 7.6, 140 mM NaCl, 1 mM EDTA, 10% glycerol, 0.5% NP-40, 0.25% Triton X-100) followed by pelleting, and 10 min rotation in 5 ml lysis buffer 2 (200 mM NaCl, 1 mM EDTA, 0.5 mM EGTA, 10 mM Tris, pH 8). Nuclei were pelleted, then swelled in lysis buffer 3 (10 mM Tris, pH 8, 1 mM EDTA, 0.5 mM EGTA, 100 mM NaCl, 0.1% deoxycholic acid, 10% N-lauryl sarcosine) for 10 min, then sonicated on high setting (30 sec on, 30 sec off) for 15 minutes to shear chromatin to less than 500 bp with Bioruptor (Diagenode). After pelleting the insoluble fraction, the supernatant was pre-cleared with Protein G agarose beads (Sigma) and anti-rabbit IgG (Sigma). 50 μg sheared chromatin was incubated in RIPA buffer (50 mM Tris, pH8, 150 mM NaCl, 1% NP-40, 0.5% sodium deoxycholate, 0.1% SDS) with either 5 μg rabbit IgG (Sigma), 5 μg mouse IgG (Santa Cruz) NELFA (Santa Cruz), 5μg H3K27ac (Abcam) or 5 μg H3K4me1 (Abcam) and recovered with Protein G agarose beads. Beads were washed twice with low salt RIPA (150 mM NaCl) and twice in high salt RIPA (300 mM NaCl), then eluted in 100 μL 1% SDS plus 100 mM sodium bicarbonate. After cross-link reversal, DNA was purified with a QIAquick PCR Purification Kit (Qiagen) and quantified by qPCR or by sequencing. ChIP-seq libraries were prepared using the Ovation Ultralow Library system (NuGen). Libraries were sequenced on the HiSeq 2500 instrument (Illumina) and ~40 million 100-nt reads were generated per ChIP sample. Sequence pre-processing and alignment were performed as described for PRO-seq.

### Coexpression analysis

Entrez gene IDs for APOE (348), APOC1 (341) and TOMM40 (10452) were submitted to COEXPRESSdb and expression data were plotted under default settings. Pearson correlation coefficients are reported.

### Expression analysis

Total RNA was isolated using the RNeasy Mini-Kit (Qiagen) and 0.5 μg RNA converted to cDNA using Taqman RT reagents kit (Thermofisher) with random hexamer priming. Gene expression was determined by SYBR green qPCR on an ABI 7900HT or BioRad CFX384 instrument using the delta-Ct method. Gene expression primers are listed in Supplemental Table 1.

### RNA abasic site detection and quantitation with ARP

RNA samples were incubated in 2 mM ARP (N-(aminooxyacetyl)-N’-(D-Biotinoyl) hydrazine) (Thermo Fisher Scientific) in 20 mM Tris-HCl, 1 mM DTT, 1 mM EDTA, pH 8.0 at 37°C with agitation for 1 h. Formaldehyde was then added to 50 mM and incubated for 10 min at 37°C to quench ARP. RNA was precipitated in 0.3 M NaAc (pH 5.5) and 3 volumes of 100% EtOH, followed by 75% EtOH washes. The samples were analyzed on 1% formaldehyde agarose gel in 1X MOPS buffer (20 mM MOPS, 5 mM Sodium acetate, 1 mM EDTA, pH 7.0). RNA was transferred to Hybond N+ nylon membrane by overnight capillary transfer in 10X SSC buffer (1.5 M NaCl, 150 mM sodium citrate, pH 7.0). Biotin signal on nylon membrane was detected using a streptavidin-based chemiluminescent method (Thermo Fisher Scientific).

### DNA-RNA hybrid immunoprecipitation (DRIP)

Immunoprecipitation procedure was adapted from previous studies (Skourti-Stathaki et al., 2011). 5×10^6^ primary fibroblasts were lysed in 600 μl cell lysis buffer (50 mM PIPES, pH 8.0, 100 mM KCl, 0.5% NP-40) and nuclei were collected by centrifugation. Pelleted nuclei were resuspended in 300 μl nuclear lysis buffer (25 mM Tris-HCl, pH 8.0, 1% SDS, 5mM EDTA). Genomic DNA, along with R-loop, were then extracted by phenol:chloroform and ethanol precipitation. Purified DNA was resuspended in IP dilution buffer (16.7 mM Tris-HCl, pH 8.0, 1 mM EDTA, 0.01% SDS, 1% Triton-X100, 167 mM NaCl) and sonicated 15 minutes in Bioruptor (Hi setting, 30sec on/30sec off) to fragments with average size of 500 bp. Three μg of S9.6 monoclonal antibody (gift from Dr. Stephen H. Leppla at NIH) or non-specific mouse IgG (Santa Cruz) was used for each immunoprecipitation. Input and precipitates were analyzed by quantitative PCR using primers (Table S2) or by sequencing.

Sequencing libraries were prepared from input and DRIP DNA using Ovation Ultralow System (NuGen) and sequenced on an HiSeq 2500 (Illumina). An average of 100 million 100 nt reads per sample were generated. Sequencing reads were pre-processed to remove adapter sequence from the end of reads using the program fastx_clipper from FASTX-Toolkit (Hannon Lab). Low-quality sequence at ends of reads represented by stretches of “#” in the quality score string in FASTQ file were also removed. Reads shorter than 35 nt after trimming were excluded from analysis. Sequencing reads were then aligned to human reference hg18 using GSNAP (Version 2013-10-28) (Wu and Nacu, 2010) using the following parameters: Mismatches ≤ [(read length+2)/12-2]; Mapping score ≥ 20; Soft-clipping on (-trim-mismatch-score=-3). Reads with identical sequence were compressed into one unique sequence. BigWig tracks were computed using bedtools and converted to hg19 coordinates using CrossMap.

### iPSC hepatocyte differentiation

Hepatic differentiation of human pluripotent stem cells was performed following a three-step protocol (Carpentier et al., 2014). First, iPSC at 60-70% confluence were treated for 3 days with 100 ng/mL Activin A and 100 ng/mL bFGF (R&D Systems) in the presence of increasing levels of FBS (0% on day 1, 0.2% on day 2, and 2% on day 3) to generate definitive endoderm. Confluent definitive endoderm cells are then passed 1:3 in presence of Rock Inhibitor on growth factors reduced Matrigel (BD Biosciences) and cultured for 8 days in differentiation medium (DMEM F12, 10% KOSR, with 1% NEAA and 1% Glutamine) containing 100 ng/mL of HGF (Peprotech) and 1% DMSO (Sigma Aldrich), to promote hepatic specification. Finally, the hepatoblasts were matured in DMSO-free differentiation medium with 10^-7^M of Dexamethasone for 3 days. Hepatocytes were then maintained for up to 1 week in hepatocyte culture medium (L15 medium, 8.4% FCS + 1% glutamine + 10% tryptose phosphate, Life Technologies) containing 1 μM insulin, 10 μM hydrocortisone and 10^-7^ M of dexamethasone (Sigma Aldrich).

### MPG Electrophoretic mobility shift assay

RNA-DNA hybrid substrates (see Table S3) containing a single N6-methyl-adenosine within the RNA strand was incubated with increasing concentration of human recombinant MPG. MPG protein was purified as described previously (Adhikari et al., 2008). Binding incubations were performed on ice for 15 min in buffer containing 20 mM Tris-HCl, pH 8.8, 10 mM (NH_4_)_2_SO_4_, 10 mM KCl, 2 mM MgSO_4_, 0.1% Triton X-100 and a range of 0-800 nM of MPG. The binding mixtures were immediately subjected to non-denaturing 6% polyacrylamide gel (acrylamide:bis-acrylamide, 37.5:1) electrophoresis. To help maintain integrity of bound complexes during electrophoresis, the gel was run at 4°C. Because some of the bound signal was, in some cases, a smear rather than a distinct band, the toolbox setting in IQTL v.8.1 was used to determine the signal more accurately in the smear after background subtraction from a box of the same size. The fraction bound relative to the total signal was then determined and plotted using Kaleidagraph v.4.1. The data were fitted to a modified Hill binding equation, where the fraction of substrate bound is related to the *Kd* as described (Ryder et al., 2008):

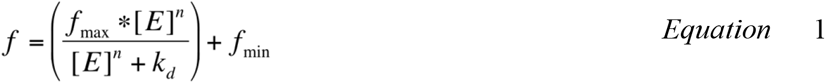

where f_max_ and f_min_ are normalization factors that represent the fraction of substrate bound at the highest and lowest asymptotes of the titration, [E] is the total enzyme concentration, and n is the Hill coefficient. The Hill coefficient measures cooperativity of binding. Values larger than n=1 may indicate positive cooperativity of multiple proteins or binding events that have not reached equilibrium. From the fitted data, the estimated *K_d_* value from triplicate binding experiments is 58 ± 16 nM.

### MPG enzymatic activity assay

RNA-DNA hybrid substrates (see Table S3) containing a single N6-methyl-adenosine were formed by heating to 90°C for 1 minute and slowly cooling to 4°C in a buffer containing 30 mM Tris, pH 7.5 and 100 mM potassium acetate. The strand containing the methylated adenosine was 5’-end labeled with fluorescein (FAM). Annealed substrates (200 nM) were incubated with (+) or without (-) 20 units of MPG in a reaction buffer containing 20 mM Tris-HCl, pH 8.8, 10 mM (NH_4_)_2_SO_4_, 10 mM KCl, 2 mM MgSO_4_, 0.1% Triton X-100 for 60 min at 37°C. Substrates were then treated with APE1 (1 μM) for an additional 2 min at 37°C. Enzymes were subsequently inactivated by a 5-minute incubation at 75°C. Samples were then denatured with formamide (1:1 vol/vol) and a 2-minute incubation at 95 °C before loading onto a 7M urea 15% denaturing polyacrylamide gel.

### RNA-sequencing

From cells treated with siMPG and siNTC, sequencing libraries were prepared using TruSeq Stranded Total RNA Library Prep Kit (Illumina). Sequencing was performed on Illumina HiSeq 2500 and > 150 million 100-nt reads were generated from each sample. Low-quality bases were trimmed from the 3’ end of reads and 3’ adaptor was trimmed using FASTQ/A Clipper with default settings (Hannon lab). Reads shorter than 35 bp were excluded from analysis. Sequencing reads were aligned to human reference (HG18) using GSNAP (Version 2013-10-28) (Wu and Nacu, 2010) using the following parameters: Mismatches % [(read length+2)/12-2];Mapping score R 20; Soft-clipping on (-trim-mismatch-score = 3)

### Precision Run-On Sequencing (PRO-seq)

PRO-seq libraries were prepared as described previously (Kwak et al., 2013; Watts et al., 2019). Fibroblast nuclei (5×10^6^) nuclei were added to 2X Nuclear Run-On (NRO) reaction mixture (final concentrations: 10 mM Tris-HCl pH 8.0, 300 mM KCl, 1% Sarkosyl, 5 mM MgCl2, 1 mM DTT, 0.03 mM each of biotin-11-A/C/G/UTP (Perkin-Elmer), 0.8 u/μl RNase inhibitor) and incubated for 3 min at 37°C. Nascent RNA was extracted by phenol (Trizol LS)/chloroform and then fragmented by base hydrolysis in 0.2 N NaOH on ice for 15 min. The reaction was neutralized by adding 0.7 X volume of 1 M Tris-HCl pH 6.8. The fragmented nascent RNA was purified using 30 μl of Streptavidin M-280 magnetic beads (Thermo Fisher Scientific) and ligated with 3’ RNA adapter (5’p-GAUCGUCGGACUGUAGAACUCUGAAC-/3InvdT/). Biotin-labeled products were recovered by streptavidin beads. The RNA products were successively treated with 5’ pyrophosphohydrolase (NEB) and polynucleotide kinase (NEB) to repair the 5’ end. RNA was ligated to the 5’ RNA adaptor (5’-CCUUGGCACCCGAGAAUUCCA-3’). The products were further purified by the streptavidin beads. RNA was reverse transcribed using RT primer (5’-AATGATACGGCGACCACCGAGATCTACACGTTCAGAGTTCTACAGTCCGA-3’). The product was PCR amplified, the resulting amplicons that are between 150-250 bp (insert >70 bp) were purified using the BluePippin (Sage Science) agarose gel electrophoresis, and then sequenced on the HiSeq 2500 instrument (Illumina) to a depth of >150 million reads per sample. PRO-seq data were aligned to the GRCh37 (hg18) build of the human genome using GSNAP (version 2013-10-28) (Wu and Nacu, 2010). BAM files were generated and normalized to reads per million mapped reads (RPM). For comparison between conditions, RPM normalized signal was plotted for AANCR-APOE locus.

The pausing index calculated using the ratio of the PRO-seq read density in the promoter to that in the gene body. In the first 1 kb of AANCR the 50 bp interval with the greatest number of reads was considered as the promoter. The gene body is the region of AANCR beyond the first kilobase.

### RNA Immunoprecipitation

Primary human fibroblasts (5×10^6^ cells per experiment) were treated with lysis buffer, (10 mM Tris-HCl pH 7.4, 10 mM NaCl, 0.5 % NP-40, 1 mM DTT, 200 units/ml RNase OUT, and EDTA-free Protease Inhibitor Cocktail), the lysate mixed with a freshly made preparation of protein A/G magnetic agarose beads bound to either anti-MPG antibody, anti m6A antibody, or anti-METTLE3 antibody. Protein-RNA complexes in the cell lysates were allowed to bind to their respective antibody-bead preparation, followed by treating the immunoprecipitated with proteinase K, RNA extraction, and DNase treatment of the extracted RNA. Samples were converted to cDNA using random hexamers and enrichment detected by qPCR.

### siRNA knockdown

Primary fibroblasts were seeded at 2 X 10^5^ per well in 6-well dishes. Cells were transfected with siRNA using Lipofectamine RNAiMax to final concentration of 12 nM on day 0 or day 3. Cells were harvested for expression or protein analysis 3 or 7 days post-transfection. Catalogue numbers for siRNA targeting MPG, METTL3 and control siRNA are listed in the key reagents table.

### S9.6 Dot Blot

S9.6 dot blot to assess genome-wide R-loop abundance was carried as before (Ramirez et al., 2021). Briefly, genomic DNA containing R-loops was incubated with 1μl of RNase H1 or mock digestion in 1X RNase H reaction buffer (10 mM Tris-HCl pH 8.0, 50 mM NaCl, 10 mM MgCl_2_, 10 mM DTT) at 37°C for 30 minutes. DNA was phenol extracted, ethanol precipitated, and reconstituted in 10 μl TE buffer. 5 μl DNA solution was loaded onto Hybond N+ nylon membranes (GE Life Sciences) presoaked with PBS, and crosslinked in UV Stratalinker 2400 (Stratagene) at the “Auto Crosslink” setting (1200 μJoulesX100). The membrane was blocked in 5% milk in PBS-0.1% Tween-20 for one hour and incubated with 1:1000 S9.6 antibody overnight at 4°C to detect RNA-DNA hybrids. A duplicate blot was incubated with anti-dsDNA antibody (Abcam) as loading control. Signal was then detected by horse-radish peroxidase (HRP)-conjugated secondary antibody and enhanced chemiluminescent, ECL, reagents. S9.6 signal normalized to dsDNA were plotted.

### Secreted APOE detection

To measure secreted APOE, cell culture media was concentrated 10-fold using an Amicon ultra centrifugal device (Sigma). Concentrated media (500 μL) was mixed with an equal volume of 2X RIPA buffer with protease inhibitor (Sigma) and PMSF (Sigma) and rotated overnight with 10 μg APOE antibody (Sigma) or IgG (Sigma). Antibody-protein complexes were recovered with protein A/G beads (Fisher) after 2 hours of rotation at 4C and three washes in 1X RIPA. Bound protein was released in 30 μL sample loading buffer, and western blot was performed by standard procedure. APOE was detected with 1:1000 primary antibody (Abcam) and 1:5000 secondary antibody. Ponceau (Sigma) staining of the western membrane was used as the loading control.

### White blood cell isolation

WBC were obtained from a subject who received clinical evaluations at the National Institutes of Health (NIH) in Bethesda, MD under IRB-approved protocol 00-N-0043 ‘‘Clinical and Molecular Manifestations of Inherited Neurological Disorders.’’ Written informed consent was received from the participant before inclusion in the study. Venous blood was collected in a 10mL lavender-top K2EDTA tube. 30mL of RBC lysis solution (Qiagen) was added to 10mL of whole blood and mixed by inverting 10 times. The sample was incubated for 5 minutes at room temperature. WBCs were pelleted by centrifugation for 2 minutes at 2,000 x *g*.

### Genetic analysis

Association of rs449647 and rs405509 with Alzheimer’s disease was analyzed using the GWAS Catalog and the NIA Genomics of Alzheimer’s Disease using the two SNP IDs individually in the search field, https://www.ebi.ac.uk/gwas/ and https://www.niagads.org/genomics/home.jsp, represented results shown in Table S1A. Genetic analysis of *APOE* expression was carried out using the eQTL calculator in GTEx (https://gtexportal.org/home/testyourown), we tested the allelic association of two SNPs in AANCR, rs449647 and rs405509 with *APOE* expression in the brain, colon, liver, colon and testis, each SNP and expression pair tested individually. The nominal P-values (<0.05) are shown in Table S1B. Allelic association of rs449647 with APOE plasma level is obtained from the publication (Rasmussen et al., 2018).

#### Key Resources Table

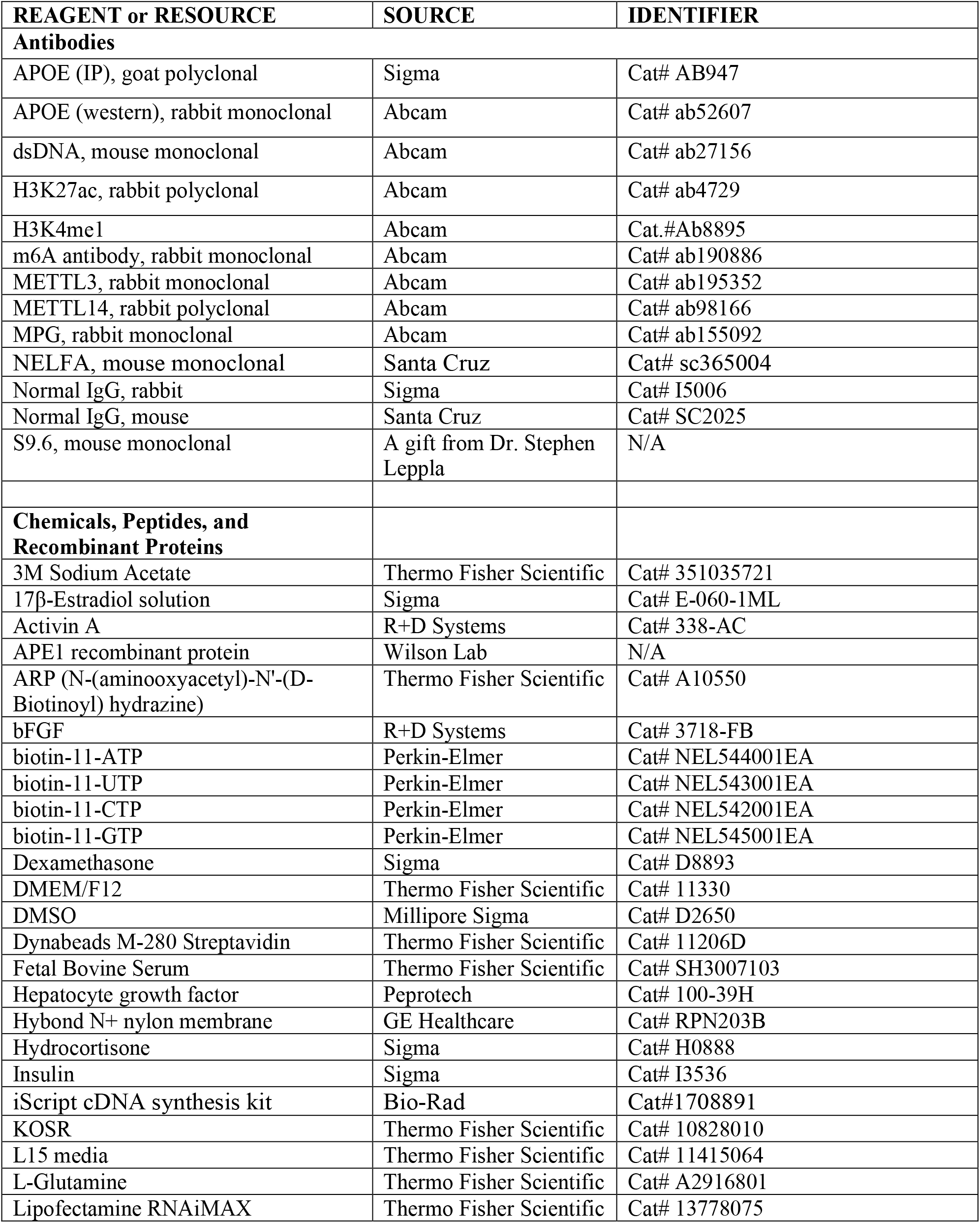
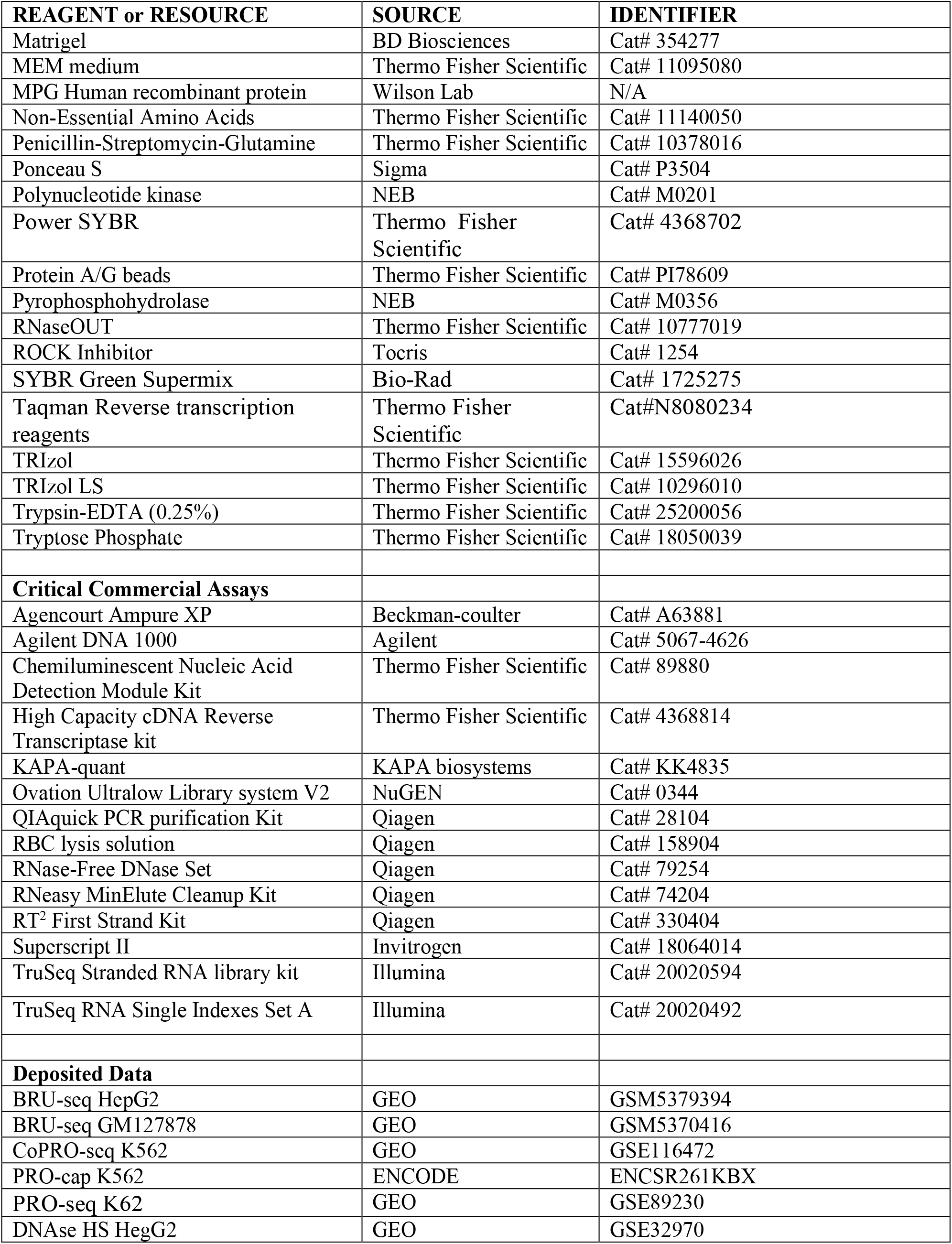
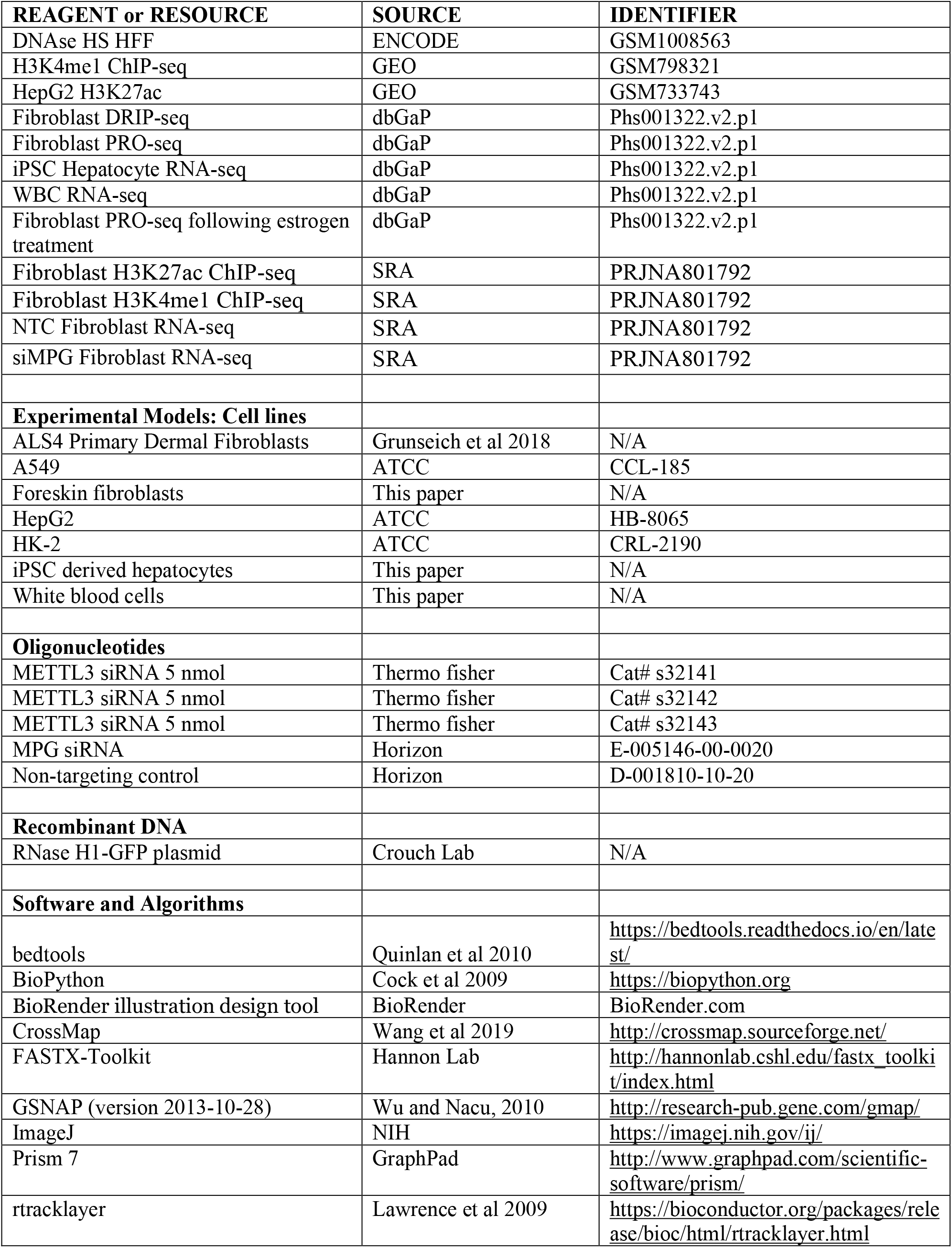
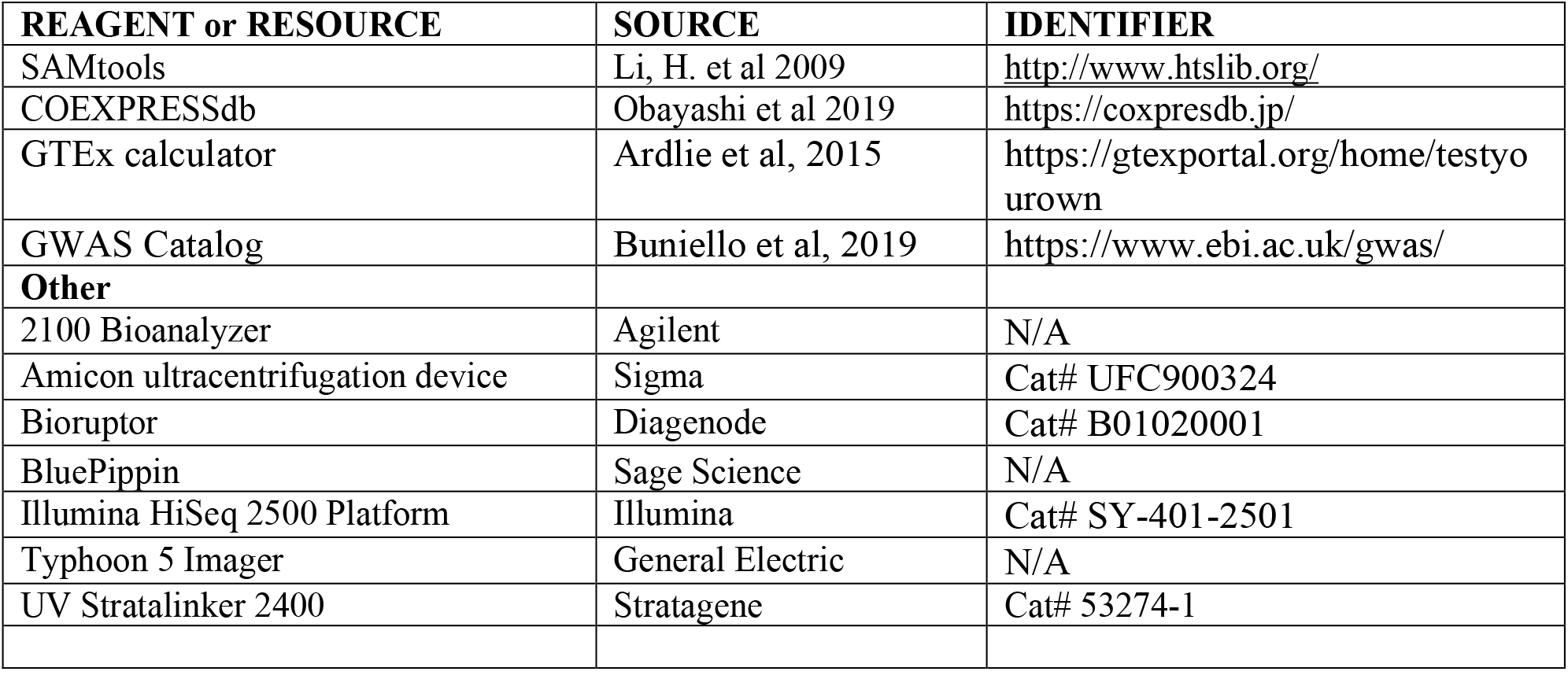

### Contact for Reagent and Resource Sharing

Further information and requests for resources and reagents should be directed to and will be fulfilled by the Lead Contact, Vivian G. Cheung.

